# Stabilization of the SARS-CoV-2 Receptor Binding Domain by Protein Core Redesign and Deep Mutational Scanning

**DOI:** 10.1101/2021.11.22.469552

**Authors:** Alison C. Leonard, Jonathan J. Weinstein, Paul J. Steiner, Annette H. Erbse, Sarel J. Fleishman, Timothy A. Whitehead

## Abstract

Stabilizing antigenic proteins as vaccine immunogens or diagnostic reagents is a stringent case of protein engineering and design as the exterior surface must maintain recognition by receptor(s) and antigen—specific antibodies at multiple distinct epitopes. This is a challenge, as stability-enhancing mutations must be focused on the protein core, whereas successful computational stabilization algorithms typically select mutations at solvent-facing positions. In this study we report the stabilization of SARS-CoV-2 Wuhan Hu-1 Spike receptor binding domain (S RBD) using a combination of deep mutational scanning and computational design, including the FuncLib algorithm. Our most successful design encodes I358F, Y365W, T430I, and I513L RBD mutations, maintains recognition by the receptor ACE2 and a panel of different anti-RBD monoclonal antibodies, is between 1-2°C more thermally stable than the original RBD using a thermal shift assay, and is less proteolytically sensitive to chymotrypsin and thermolysin than the original RBD. Our approach could be applied to the computational stabilization of a wide range of proteins without requiring detailed knowledge of active sites or binding epitopes, particularly powerful for cases when there are multiple or unknown binding sites.

## Introduction

Many natural proteins are marginally stable (Goldenzweig and Fleishman 2018, and improving their stability is a common prerequisite for diverse industrial and medical applications, from engineering thermostable proteases for laundry detergents (Zhao and Arnold 1999; Wintrode et al. 2000) to immunogen design like the pre-fusion stabilizing coronavirus Spike protein mutations used in several regulatory approved COVID vaccines (Pallesen et al. 2017, Baden et al. 2021, Mulligan et al. 2020). However, many stabilizing mutations can reduce activity or function in the engineered protein (Beadle and Shoichet 2002). Thus, balancing the tradeoff between improved stability and maintaining function remains a major challenge for the protein designer. Previous work to predict non-disruptive stabilizing mutations have incorporated factors such as evolutionary conservation (Goldenzweig et al. 2016), distance to an active site or binding site (Tokuriki et al. 2007), and the local packing density (Klesmith et al. 2017, Wrenbeck et al. 2019). These and other prediction methods can successfully identify stabilizing mutations with neutral impact on activity without large directed evolution campaigns.

However, such prediction methods identify potential stabilizing mutations that are located predominantly at or peripheral to the solvent-contacting surface of the protein. Additionally, the set of mutations identified typically are not in direct contact and thus are usually independent of other mutations (Goldenzweig et al. 2016). Yet for many applications, like immunogen or diagnostic reagent design, the surface of the protein must remain unchanged. Design strategies then would be akin to remodeling a historic building, in which the interior may be reinforced and updated as needed within the requirement that the exterior structure is preserved intact. In this stringent design case, the surface structure can be held in place during computational design with constraints on movement, while interior, non-solvent-exposed residues are allowed to vary as necessary barring major deformation of the protein backbone. Stabilizing protein vaccine immunogens is an example of a relevant application because effective stabilized antigens must retain the ability to bind many different antibodies at distinct epitopes across the protein surface and only a small fraction of the epitopes are known. Rather than risk damaging these epitopes, protein core redesign can be used on the protein from the inside out without making significant changes to the surface structure.

One such potential protein of immediate global importance is the SARS-CoV-2 receptor binding domain (RBD), a diagnostic antigen (Premkumar et al. 2020) and vaccine immunogen (Tai et al. 2020, Chen et al. 2017). RBD is critically important as the titer of anti-RBD antibodies are the major correlates of protection in SARS-CoV-2 (Yu et al. 2020, Feng et al. 2021, Rogers et al. 2020). The RBD is a 205 amino acid domain of the SARS-CoV-2 Spike protein that contains distinct epitopes for anti-Spike neutralizing antibodies (Francino-Urdaniz et al. 2021; Dejnirattisai et al. 2021, Barnes et al. 2020). The protein structure is composed primarily of beta sheets with connecting loop and helices and contains nine cysteine residues, eight of which form four disulfide bonds: C336-C361, C379-C432, C480-C488, and C391-C525 (Lan et al. 2020) (**Fig 1A**). The receptor binding motif (RBM) spans residues 438-506 and contains the binding surface for the cell receptor ACE2 (Lan et al. 2020, Wrapp et al. 2020). Away from the RBM, the protein core contains several under-packed regions, most notably a cryptic linoleic acid binding pocket surrounded by positions F338, V341, F342, I358, C361, A363, L368, F374, C379, L387, and F392 (Toelzer et al. 2020) and a buried unsatisfied polar group at Y365. The presence of under-packed hollow cavities and side chains with unsatisfied hydrogen bonds in the core suggests that mutations could improve stability, as successfully demonstrated by the introduction of key RBD mutations into this fatty acid binding pocket by the King lab (Ellis et al. 2021). In a different approach, the King lab also reduced protein aggregation by mutating hydrophobic patches on the surface (Dalvie et al. 2021), though this could potentially remove critical epitopes recognized by the immune system.

**Figure 1:**
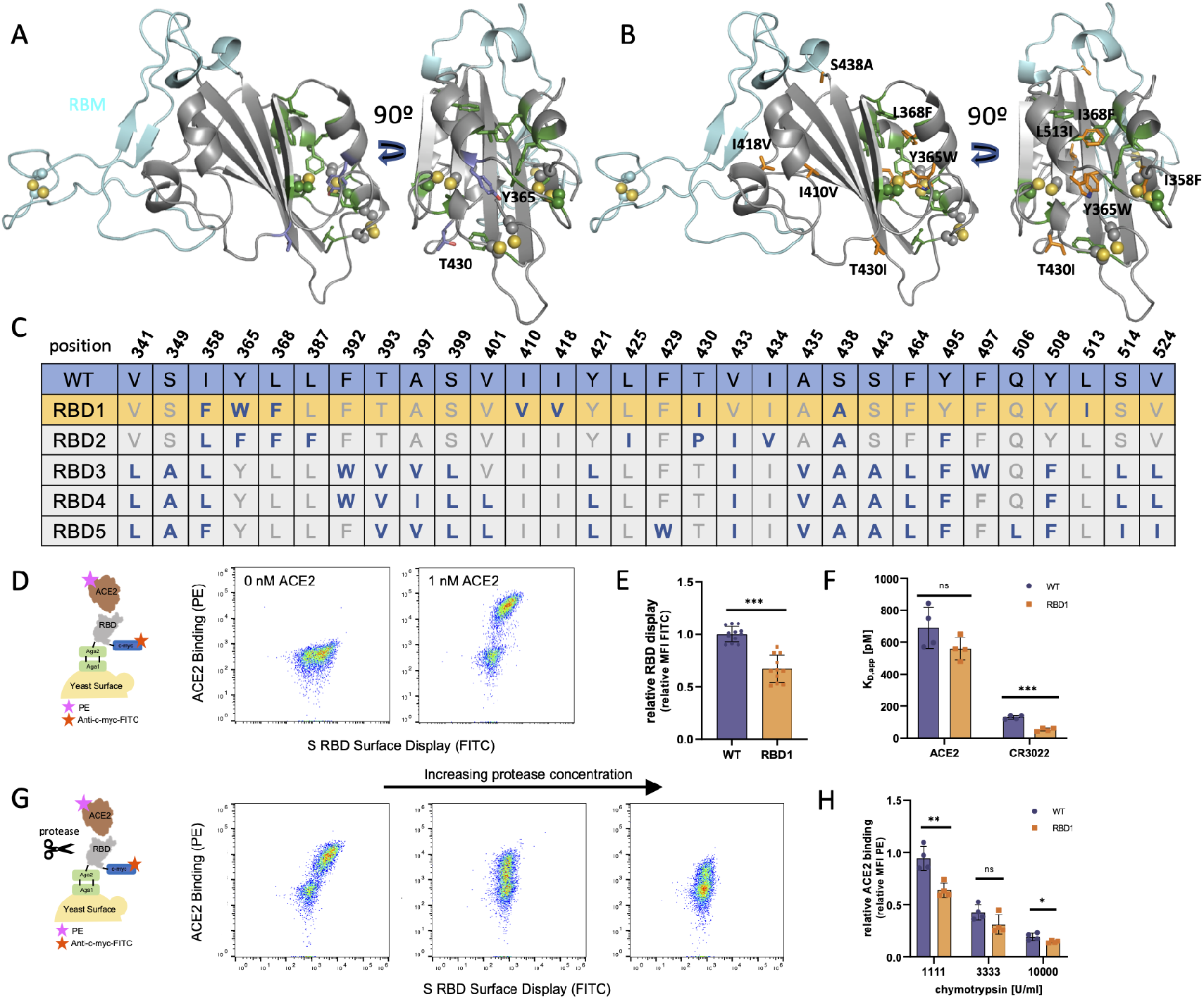
Evaluation of SARS-CoV-2 S RBD computational designs using yeast display. **A**. Structure of SARS-CoV-2 S RBD (PDB ID 6M0J). The receptor binding motif (RBM) for ACE2 spanning positions 438-506 is colored in cyan, residues adjacent to cryptic fatty acid binding pocket colored green, disulfide bonds shown as spheres and colored by element. Residues T430 and Y365 containing buried unsatisfied hydrogens colored in blue. **B**. RBD structure with sequence differences for the RBD1 design shown as orange sticks. **C**. Sequence diversity of the five RBD designs. **D**. Cartoon of yeast surface display assay and representative cytograms of yeast displayed WT RBD in the presence and absence of biotinylated ACE2 at the indicated concentrations. **E**. Relative RBD surface expression of WT (n=12) and RBD1 (n=11) as measured by relative FITC mean fluorescence intensity using an anti-myc FITC antibody (p-value = 2e-7). **F**. Apparent K_D_ (K_D, app_) for the interactions between ACE2 (p-value = 0.13) and CR3022 (p-value = 1.2e-4) and surface-displayed WT and RBD1 (n=4). **G**. Cartoon and cytograms showing protease susceptibility assay for protein stability in yeast surface display. Displayed RBD is treated with protease (chymotrypsin) to denature the RBD before binding with ACE2. Treatment with higher concentrations of protease will result in more highly denatured RBD protein that is less capable of binding ACE, leading to lower PE signal. **H**. Relative binding signal for WT and RBD1 after treatment with 1111, 3333, and 10000 U/ml chymotrypsin, measured by relative PE mean fluorescence intensity (p-value = 0.0039 at 1,111 U/ml, p-value = 0.096 at 3,333 U/ml, p-value = 0.039 at 10,000 U/ml chymotrypsin, n=4).

In this work, we present the stabilization of RBD by mutating key surface epitopes and using computational design and deep mutational scanning to identify stabilizing core mutations on RBD. Combining a subset of these mutations led to at least one improved RBD design with greater comparative proteolytic and thermal stability while maintaining recognition by a panel of antibodies binding multiple distinct epitopes. These designs may be useful as diagnostic and vaccine immunogens. More fundamentally, our work advances our understanding of critical factors influencing the successful computational redesign of cores of existing proteins.

## Materials and Methods

### Plasmid constructs

All plasmids and primers used for this work are listed in **Tables S1 and S2**, respectively. All plasmids were verified by Sanger sequencing. The yeast display plasmid pETconV4 (production_PETCON_V4_B1A2), optimized for use in the protease susceptibility assay, was received as a gift from the University of Washington (Maguire et al. 2021). pJS699 (Wuhan-Hu-1 S-RBD(333-537)-N343Q for fusion to the C-terminus of AGA2) was previously described (Banach et al. 2021).

To construct plasmids pACL002-pACL006 and pACL009-pACL015 containing all RBD design sequences for yeast surface display, DNA sequences were ordered as gBlocks (IDT) and cloned into pETconV4 using restriction enzymes NdeI and XhoI (NEB).

To construct plasmid pACL007, the wild-type RBD sequence was amplified from plasmid pJS699 using primers forward_pJS699_RBD_pETconV4 and reverse_pJS699_RBD_pETconV4 to add regions of pETconV4 homology surrounding the RBD sequence. Plasmid pETconV4 was linearized by digestion with NdeI and XhoI (NEB). Both fragments were run on a 1% agarose gel and purified using a Monarch DNA Gel Extraction kit (NEB). The fragments were then joined using the NEBuilder HiFi DNA Assembly protocol (NEB).

DNA sequences for plasmids pACL002-pACL006 and Design1_pETconV4-Design7_pETconV4 containing all RBD design sequences were ordered as gBlocks (IDT) and cloned into pETconV4 using restriction enzymes NdeI and XhoI (NEB).

Plasmid pACL008 was created by inserting the BbvCI restriction enzyme site into the pACL005 plasmid using site-directed mutagenesis. PCR primers add_BbvCI_forw_pETconV4 and add_BbvCI_rev_pETconV4 were used to insert the BbvCI restriction enzyme site using KAPA HiFi HotStart ReadyMix (Roche). The PCR product was run on a 1% agarose gel and extracted to purify. 1 µL of the purified product was ligated into a circular plasmid using KLD reaction mix (NEB) incubated for 10 min at room temperature.

RBD designs to be tested further as soluble proteins were codon optimized for *P. pastoris* (IDT), ordered as gBlocks, and cloned into the pPICZ*α*A secreted expression vector (ThermoFisher V19520) using EcoRI and SacII restriction sites. RBD designs in *P. pastoris* do contain the N343 glycan that was removed from designs optimized for yeast surface display via an N343Q mutation. N343 was reintroduced to RBD sequences cloned into *P. pastoris* plasmids by Q5 site-directed mutagenesis (NEB) using primers Q343N_SDM_WT_D1_D3_F and Q343N_SDM_R.

### Computational Protein Design

Initial designs were generated using two separate Rosetta-based methods (Leman et al. 2020). RBD1 and RBD2 were designed using FuncLib (Khersonsky et al. 2018) run on PDB entry 6VSB, using a PSSM threshold of 0 and ΔΔG of 3 Rosetta energy units. The RBD was manually split to two spatial sub-domains to reduce the combinatorial complexity. Positions in sub-domain 1: 341, 342, 358, 365, 368, 377, 387, 392, 395, 397, 431, 434, 511, 513, 515, 524. Positions in sub-domain 2: 350, 398, 400, 401, 402, 410, 418, 419, 423, 425, 430, 433, 438, 442, 495, 497, 507, 510, 512. FuncLib results for each sub-domain were sorted by Rosetta energy, and the best designs separated by at least three mutations from one another were chosen. All chosen mutants from both sub-domains were combined, modeled and ranked by energy. The resulting combined designs were clustered as described above. The 30 best scoring combined designs were manually inspected, and RBD1 and 2 were chosen for experimental testing.

Proteins RBD3, RBD4, and RBD5 were designed using FastDesign (Maguire et al. 2021) (access date 15-April-2020) on an RBD structure (PDB ID: 6MOJ) prepacked using FastRelax, with alternating cycles of repacking and minimization with design. Instead of fixing the backbone, coordinate constraints were applied to non-core residues, with constraints scaled to the B-factor of that atom. Core residues were identified by the layer selection command, with the exclusion of cysteines.

To introduce additional mutational variation into designs RBD7-12, additional rounds of Rosetta design were performed using different RBD structures (PDB ID: 6M0J, 7JMO, and 6LZG), as well as varying scaling of B-factor constraints to allow more or less flex to the protein surface.

Allowed mutations were selected using a resfile in which the default was NATAA and residues with high expression identified in the deep mutational scanning experiment described here and/or in Starr et al (Starr et al. 2020) were allowed to mutate to the best residue using PIKAA to either the wild-type identity or any of the possible beneficial mutations.

### Recombinant protein production, purification, and preparation

ACE2-Fc, produced and purified following Walls et al. 2020 (Walls et al. 2020), and CR3022 (ter Meulen et al. 2006) were kind gifts from Neil King’s lab at the University of Washington. The anti-SARS-CoV-2 RBD antibody panel used (CC6.29, CC6.32, CC6.33, CC12.1, CC12.7) was a kind gift from Dennis Burton’s lab at Scripps and were produced and purified according to Rogers et al. (Rogers et al. 2020). Background binding level in ELISA assays was measured using Human IgG Isotype Control (ThermoFisher #02-7102).

RBD designs were produced recombinantly in *Pichia pastoris* as follows. pPICZ*α* vectors (ThermoFisher V19520) containing WT RBD or RBD designs were linearized by SacI and greater than 5 *μ*l were transformed into electrocompetent *P. pastoris* X-33 (ThermoFisher C18000) at 2000V using a 2 mm electroporation cuvette (Bulldog Bio) and Eppendorf electroporator and then plated on yeast extract peptone dextrose plus sorbitol plates (YPDS: 1% w/v yeast extract, 2% w/v peptone, 2% v/v glucose, plus 1.0M sorbitol) supplemented with 100 *μ*g/ml zeocin (ThermoFisher 25001). Single colonies were selected and streaked on Minimal Dextrose (1.34% w/v yeast nitrogen base, 4×10^−5^% w/v biotin, 2% v/v glucose), Minimal Methanol (1.34% w/v yeast nitrogen base, 4×10^−5^% w/v biotin, 0.5% v/v methanol), and YPDS plus 400 *μ*g/ml zeocin to identify the colonies with MutS phenotype that are most resistant to zeocin. Integration of the RBD sequence was confirmed using colony PCR using primer colonyPCR_pichiaRBD_rev.

For RBD production, single colonies were used to inoculate buffered complex glycerol (BMGY) medium (1% w/v yeast extract, 2% w/v peptone, 1.34% w/v YNB, 400 *μ*g/L biotin, 0.1 M potassium phosphate, pH 6.0, 1% v/v glycerol), scaling up to 1 L volume grown at 30°C with 250 rpm agitation until the culture reached OD_600_=2-6. Cells were harvested, resuspended in buffered methanol complex medium (BMMY: 1% w/v yeast extract, 2% w/v peptone, 1.34% v/v 10X yeast nitrogen base, 400 *μ*g/L biotin, 0.1 M potassium phosphate, pH 6.0, 0.5% v/v methanol), and incubated at 30°C with 250 rpm agitation for 4 days, adding 0.5% v/v methanol daily to maintain induction. Cells were removed by centrifugation at 3200 x g for 10 min.

RBD harvest and Ni^2+^-NTA column purification was done as described (Argentinian AntiCovid Consortium 2020). After elution with imidazole the purified protein was concentrated using a 10 kDa MW cutoff Amicon centrifugal filter (Sigma), buffer exchanged into PBS (10mM Na_2_HPO_4_, 1.8 mM KH_2_PO_4_, 2.7mM KCl, 137 mM NaCl, pH 7.4) using PD-10 desalting columns (GE Healthcare), and stored at 4°C. Protein was quantified by absorbance at 280 nm using the theoretical extinction coefficient derived from the protein sequence when all four disulfide bonds are intact (WT: 33850 M^-1^cm^-1^, RBD6: 37860 M^-1^cm^-1^, RBD8: 37860 M^-1^cm^-1^, RBD10: 36370 M^-1^cm^-1^). To visualize protein bands, samples were denatured for 10 min at 99°C in SDS sample buffer (188 mM Tris-Cl, 3% w/v SDS, 30% v/v glycerol, 0.01% v/v bromophenol blue) plus 100 mM DTT and separated by 4-20% gradient SDS-PAGE (BioRad 4568096). To visualize the N343 glycan, samples were incubated for 1 h at 37°C with 1 µl Endo H in GlycoBuffer 3 (NEB P0702S) after denaturing in SDS sample buffer prior to running SDS-PAGE.

### Yeast display titrations and protease susceptibility screening

For initial screening of the initial designs compared to wild type, cell surface titrations of EBY100 *S. cerevisiae* harboring the various RBD display plasmids were grown in 1ml M19D (5 g/l casamino acids, 40 g/l dextrose, 80 mM MES free acid, 50 mM citric acid, 50 mM phosphoric acid, 6.7 g/l yeast nitrogen base, adjusted to pH7 with 9M NaOH, 1M KOH) overnight at 30°C. Expression was induced by resuspending the M19D culture to OD_600_=1 in M19G (5 g/l casamino acids, 40 g/l galactose, 80 mM MES free acid, 50 mM citric acid, 50 mM phosphoric acid, 6.7 g/l yeast nitrogen base, adjusted to pH7 with 9M NaOH, 1M KOH) and growing 22 h at 22°C with shaking at 300 rpm. Yeast surface display titrations for hACE2-Fc and CR3022 IgG were performed as described by Chao et. al. (Chao et al. 2006) with an incubation time of 3h at room temperature and using secondary labels 0.6 µL anti-c-myc-FITC (Miltenyi Biotec) and 0.25 µL Goat anti-Human IgG Fc PE conjugate (ThermoFisher #12-4998-82). Titrations were performed in biological replicates (cultures from independent colonies grown on separate days) and technical replicates (n = 4). The levels of display and binding were assessed by fluorescence measurements for fluorescein (FITC) and phycoerythrin (PE) using a Sony SH800 cell sorter equipped with a 70 µm sorting chip and 488 nm laser. Cells for protease susceptibility measurements were grown and prepared for titrations as described by Chao et. al. (Chao et al. 2006). Cells were first treated with 1,000-4,000 U/mL chymotrypsin (Sigma #XC4129) for 5 min as described in Rocklin et. al. (Rocklin et al. 2017), using a 200 µL volume. Chymotrypsin activity was determined relative to trypsin using the Pierce fluorescent protease assay (ThermoFisher), and trypsin concentration was quantified using an enzymatic assay with *N*_*α*_-Benzoyl-L-arginine ethyl ester (BAEE, Sigma-Aldrich) exactly according to Rocklin et al (Rocklin et al. 2017) except with volumes scaled to be read on a 96-well plate reader. After chymotrypsin treatment, cells were incubated with 200 nM hACE2-Fc for 3h at room temperature, washed with Tris buffered saline + BSA (TBSF), and labeled with 0.6 µL anti-c-myc-FITC (Miltenyi Biotec) and 0.25 µL Goat anti-Human IgG Fc PE conjugate (ThermoFisher #12-4998-82). Protease susceptibility measurements were performed in biological replicates (n = 2) and technical replicates (n = 2).

For screening of subsequent designs, *S. cerevisiae* EBY100 harboring the various RBD display plasmids were grown from -80 °C cell stocks in 1ml SDCAA for 4-6h at 30°C. Expression was induced by resuspending the SDCAA culture to OD_600_=1 in SGCAA and growing at 22h at 22°C with shaking at 300rpm. Yeast surface display titrations and protease susceptibility measurements were performed as described above, except with an incubation time with hACE2-Fc or CR3022 of 4h at room temperature.

### Preparation and screening of mutagenic library

We identified 90 positions where the RBD1 residue has a relative solvent accessibility of less than or equal to 20%. Relative solvent accessibility was calculated by normalizing the solvent-accessible surface area of the RBD1 design calculated using dssp (Kabsch and Sander 1983) relative to the maximum theoretical solvent accessibility of each residue (Tien et al. 2013). These 90 positions were mutated to every other amino acid plus stop codon by comprehensive nicking mutagenesis (Wrenbeck et al. 2016) using NNK primers (**Table S2**) and template plasmid pACL008. For compatibility with 250 bp paired end Illumina sequencing, the mutagenic library was divided into two tiles. Tile 1 encompassed positions 336-430 and tile 2 encompassed positions 431-524. Biological replicates of each library were prepared by separate nicking mutagenesis reactions. Library plasmids were transformed into chemically competent EBY100 cells as described (Medina-Cucurella and Whitehead 2018). Yeast stocks were stored in yeast storage buffer (20% w/v glycerol, 200mM NaCl, 20mM HEPES pH 7.5) at -80°C. Serial dilutions were plated on SDCAA and incubated 3 days to calculate the transformation efficiency.

To screen the core residue mutagenic library for protease-resistant mutations, library yeast stocks for each tile and biological replicate were thawed, centrifuged for 3 min at 2500 x g, resuspended in 1 ml of SDCAA, and grown for 4-6h at 30°C. Expression was then induced by resuspending the SDCAA culture to OD_600_=1 in 1 ml SGCAA and growing at 18h at 22°C with shaking at 300rpm, after which cells were resuspended in 1 ml TBSF at OD_600_=2 (2×10^7^ cells). Cells in TBSF were treated with either 0 (reference population), 1000, 2000, or 4000 U/ml chymotrypsin as described above, incubating for 5 min at room temperature with an occasional vortex, then spun down at 2,500 x g and washed with 2ml TBSF three times. After chymotrypsin treatment, cells were incubated with 200nM hACE2-Fc in a 10:1 hACE2-Fc/displayed protein ratio (Medina-Cucurella and Whitehead 2018) for 4 h at room temperature, washed with TBSF, labeled with 50 µL Goat anti-Human IgG Fc PE conjugate (ThermoFisher Scientific Invitrogen Catalog # 12-4998-82) diluted in 1.95 mL TBSF for 10 minutes covered on ice, and washed again with TBSF. Labeled cells were sorted on a Sony SH800 cell sorter, with an SSC-A/FSC-A gate used to identify cells, a FSC-H/FSC-A gate to select only single cells, and a PE-A/FITC-A gate used to identify cells that bind hACE2-fc (**Fig S1**). For both chymotrypsin-treated populations and the no-protease reference population, cells binding hACE2-fc were collected using the PE/FITC gate with a PE fluorescence signal greater than 2000. A minimum of 170,000 cells for tile 1 and 130,000 cells for tile 2 were collected to compile at least 100-fold more cells than the theoretical sub-library size (Medina-Cucurella and Whitehead 2018). Collected cells were recovered in SDCAA with 1x PenStrep for 30h then frozen at -80 °C in yeast storage buffer in 1 mL aliquots at OD_600_=4.

### Deep sequencing preparation

Libraries were prepared for deep sequencing as described in Medina-Cucurella and Whitehead (Medina-Cucurella and Whitehead 2018), using a Zymo Yeast Plasmid Miniprep II kit (Zymo Research) and a Monarch PCR & DNA Cleanup kit (NEB) with the following changes. Inner primers RBD-F1_tile1_F_Illumina and RBD-F1_tile1_R_Illumnia were used to amplify tile 1 with an annealing temperature of 70°C; inner primers RBD-F1_tile2_F_Illumina and RBD-F1_tile2_R_Illumnia were used to amplify tile 2 with an annealing temperature of 63°C. 5 µL of PCR product from the inner primer amplification was cleaned used 0.5 µL Exonuclease I (NEB) and 1 µL rSAP (NEB), incubating for 15 min at 37°C then 15 min at 80°C. 2 µL of purified DNA was carried forward to the 2^nd^ PCR reaction. Samples were purified using Agencourt Ampure XP beads (Beckman Coulter), quantified using PicoGreen (ThermoFisher), pooled, and sequenced on an Illumina MiSeq using 2 × 250 bp paired-end reads at the BioFrontiers Sequencing Core (University of Colorado, Boulder, CO). Library statistics are listed in **Table S3**.

### Deep sequencing analysis

All deep sequencing data analysis was performed using the Protein Analysis and Classifier Toolkit (Klesmith and Hackel 2019) available at GITHUB (https://github.com/JKlesmith/PACT). Analysis was performed using the “fitness” protocol with “mutationtype: single”, “mutthreshold: 1”, “min_coverage: 0.2”, “qaverage: 20”, “ref_count_threshold: 5” and “sel_count_threshold: 5”.

### ELISA binding affinity measurements

50 µL of each purified soluble RBD variant at a protein concentration of 2 µg/ml was immobilized to Microlon clear plates (Greiner Bio-One 655081) at 4°C overnight. Plates were sealed using Microseal B adhesive sealers (BioRad MSB-1001). The following day, the solutions were decanted by flicking and all wells were washed three times with 200 µL of PBSM (PBS + 0.1% Tween-20 + 3% non-fat milk) followed by vigorous tamping on a pad of paper towels to remove residual liquid. Plates were blocked using 100 µL PBSM for 1 h at room temperature. Blocked plates were decanted by flicking. Binding reactions were assembled using serial dilutions of hACE2-Fc in PBSM from 100 pM – 12.8 nM, or appropriate concentrations of binding antibody. Plates were transferred to a plate shaker (Heidolph Titramax 1000) and incubated at room temperature for 4 h with shaking at 400 rpm. Plates were decanted by flicking and were washed three times with 200 µL PBSM followed by vigorous tamping on a pad of paper towels. Detection was enabled by addition of 100 µL Goat anti-Human IgG Fc secondary antibody conjugated to horseradish peroxidase (ThermoFisher #A18817), diluted 1:50,000 into PBSM and incubated in the above plate shaker for 1 h at room temperature. Binding was visualized through the addition of 50 µL 1-Step™ Ultra TMB-ELISA Substrate Solution (ThermoFisher, #34028), incubated in the above plate shaker for approximately 5 minutes. TMB development was quenched by addition of 50 µL of 2M sulfuric acid and plates were read at 450 nm using a Biotek Synergy H1 Hybrid multimode plate reader. The K_D,app_ for each reaction was calculated using non-linear least squares regression performed using custom Python scripts.

### Thermal melt assay

WT and RBD design apparent melting temperatures, T_m, app_, were measured using a SYPRO Orange thermal shift assay (Huynh and Partch 2015). 2µl of 200x SYPRO Orange (Life Technologies) was added to 2 µg purified RBD protein in 18 µl PBS in a MicroAmp Fast 96-Well Reaction Plate (0.1ml) (Applied Biosystems), for a total reaction volume of 20 µl. Plates were sealed with Micoseal ‘B’ seals (BioRad). Thermal melt analysis was performed in a QuantStudio 6 Flex RT-PCR device (ThermoFisher) over a range of 25-85°C with 1°C change per minute and with a 2-min incubation at the first and last temperatures. T_m,app_ was determined by calculating the minimum of the first derivative of measured fluorescence.

### In vitro proteolysis assay

The proteolytic stability of RBD designs was tested as described in Whitehead et. al. (Whitehead et al. 2009) with the following changes. 1 µg RBD protein was incubated in 50 mM Tris-HCl pH 8.0, 0.5 mM CaCl_2_ at 37°C for 1 h in the presence and absence of 0.0067 mg/ml thermolysin, for a total reaction volume of 15 µl. Reactions were inactivated with 5 µl of 50 mM EDTA, then 10 µl of the reaction was mixed with 5 µl 3x SDS loading buffer with 100 mM DTT, denatured at 99°C for 10 minutes, and run on an SDS-PAGE gel. Gel bands were quantified using ImageJ software (Abramoff et al. 2004) to determine the relative density of each band, compared to the average of 3 control (no thermolysin) samples on the same gel.

## RESULTS

We hypothesized that computational design of the protein core could improve the stability of the RBD from the original Wuhan-Hu-1 SARS-CoV-2 virus without impacting its ability to bind ACE2 or a panel of neutralizing antibodies. We used two different Rosetta design protocols (see Methods) to introduce mutations into the RBD protein core, leaving the four disulfide bonds intact. The first method used the FuncLib automated design method (Khersonsky et al. 2018) to target mutations to two distinct core regions away from the RBM, separated by an internal beta sheet. FuncLib starts with a phylogenetic analysis, restricting design at each of the selected amino acid positions only to mutations that are commonly observed among natural homologs. Then, the method models each mutation in Rosetta, allowing the structure to adapt to the mutation using whole-protein minimization and filtering mutations that are highly destabilizing. Finally, all combinations of mutations at the allowed positions are enumerated using Rosetta atomistic design, relaxed using whole-protein minimization and ranked by energy. Unlike other stability-design methods that do not relax all possible mutants (Goldenzweig et al. 2016), FuncLib finds rare combinations of stabilizing constellations of amino acids that can stabilize the core of the protein (Warszawski et al. 2019, VanDrisse et al. 2021). The resulting designs RBD1 and RBD2 mainly increase hydrophobic packing and remove buried unsatisfied hydrogen bonds like those on Y365 and T430 (**Fig 1B**), resulting in 8 and 10 mutations, respectively, from the WT RBD sequence (**Fig 1C**). The second method used the Rosetta FastDesign packing method (Maguire et al. 2021) to target all core residues while restricting surface residue movement using constraints. The resulting designs, RBD3-RBD5, were more aggressive than RBD1-2 as they contained a combination of either 18 or 19 mutations from WT spread more evenly throughout the core (**Fig 1C**). All designs concentrated on several small to large mutations in the underpacked cryptic linoleic acid binding pocket, including at V341, I358, L368, L387, and F392 (Toelzer et al. 2020) (**Fig 1B**).

The five designs were ordered as synthetic genes, subcloned into a plasmid optimized for the protease susceptibility assay (Maguire et al. 2021), and expressed using a yeast surface display platform (**Fig 1D**) previously demonstrated to display properly folded aglycosylated S RBD (Wuhan Hu-1 S RBD(333-537)-N343Q; wild-type or “WT”) (Banach et al. 2021; Francino-Urdaniz et al. 2021). Expression of the aglycosylated constructs was induced at 22°C for 22 hours in M19G media (see Methods). Protein surface display level assessed by expression of a C-terminal c-myc epitope tag showed that, of the five initial designs tested, only RBD1 displayed on the yeast surface (**Fig 1E;** data not shown), suggesting major structural changes of the other designs leading to improper folding. Additionally, RBD1 displays at a significantly lower level than WT (p-value = 2e-7, n = 11) (**Fig 1E**). This suggests that RBD1 is less stable than WT, since surface expression is known to correlate with stability (Klesmith et al. 2017).

Next, we assessed whether RBD1 maintains fidelity of critical surface epitopes by screening for maintenance of binding to both ACE2 and the antibody CR3022 (Yuan et al. 2020), which binds at a distinct epitope from ACE2. Titrations of ACE2 and CR3022 showed no significant difference at the 95% confidence level (p-value = 0.13, n = 4) in the apparent dissociation constant, K_d,app_, between the wild type sequence and RBD1 for ACE2, while the CR3022 K_d,app_ was slightly but significantly lower (p-value = 1.2e-4, n = 4) for RBD1 than WT (**Fig 1F**). Relative protein stability was assessed using a yeast surface protease susceptibility assay (Rocklin et al. 2017) in which incubation with chymotrypsin cleaves RBD, resulting in diminished ACE2 binding under saturating ACE2 concentrations (**Fig 1G**). Treatment with increasing concentrations of chymotrypsin showed that yeast displayed RBD1 is more proteolytically sensitive than WT (p-value = 0.0039 at 1,111 U/ml chymotrypsin and p-value = 0.039 at 10,000 U/ml chymotrypsin, n = 4) (**Fig 1H**). The higher proteolytic sensitivity and the lower overall surface expression of RBD1 relative to WT suggests that while RBD1 maintains the surface topology of WT, it is a comparatively less stable protein.

We hypothesized that deep mutational scanning could identify mutations that improve the stability of the RBD1 design (**Fig 2A**). We first identified all 90 positions where a RBD1 residue had a relative solvent accessibility of 20% or less. Next, a site-saturation mutagenesis library of these positions was created using nicking mutagenesis (Wrenbeck et al. 2016). The full library was split into two sub-libraries (“tiles”), with tile 1 encompassing 52 positions between positions 336-430 and tile 2 encompassing 38 positions between 431-524. Additionally, replicates (labeled “A” and “B”) for each tile were prepared, resulting in a total of four separate libraries.

**Figure 2:**
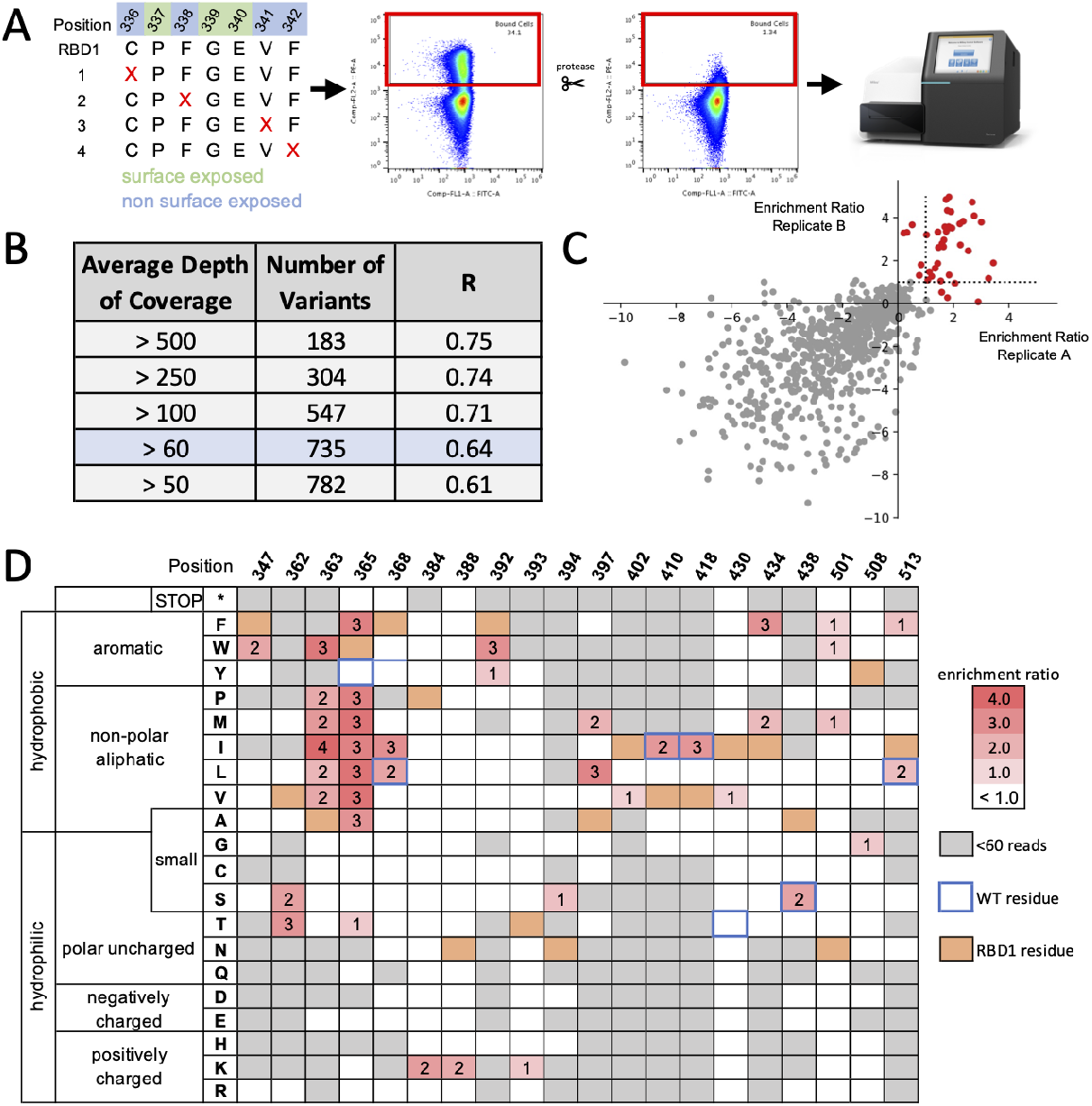
Deep mutational scanning of RBD1 using a yeast-based proteolysis assay. **A**. A flow diagram of the protease susceptibility deep mutational scanning protocol. **B**. Correlation coefficient R of enrichment ratios between biological replicates as a function of average depth of coverage. **C**. Scatterplot of enrichment ratios between replicates at an average depth of coverage of 60 or more reads. Mutations with an average enrichment ratio greater than 1.0 and positive enrichment ratios for both replicate A and B are shown as red closed circles, with all other mutations shown as closed grey circles. **D**. Heatmap of hits from the deep mutational scan. Point mutations with a read depth of less than 60 colored gray. Hits are color-coded by average enrichment ratio in shades of red. All other mutations are colored white. Blue boxes indicate positions where the WT residue is distinct from the RBD1 residue (colored orange).

Libraries were transformed into yeast (*S. cerevisiae* EBY100 (Boder and Wittrup 1997)) and induced by galactose to display RBD1 mutants. Cells were then treated with four different concentrations of chymotrypsin (a reference at 0 and samples at 1000, 2000, 4000 U/ml), and then screened for maintenance of ACE2 binding. For each sample, cells maintaining ACE2 binding were collected by fluorescence-activated cell sorting (FACS). After outgrowth, plasmid DNA was purified, prepared, and deep sequenced. Coverage of all possible single nonsynonymous mutations ranged from 71.9% to 77.7% depending on the tile and replicate. In all, 1,533/1,800 (74%) possible single point mutants at the 90 core positions were present in the screened libraries, with complete library statistics reported in **Table S3**.

We next evaluated the normalized enrichment ratio of each mutant i (ε_i_) observed in the population relative to the RBD1 sequence (ε_RBD1_) using the following equation:

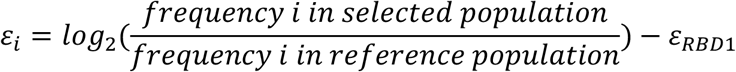

Because our reference population included a selection for maintenance of ACE2 binding, the 1,533 mutations observed by deep sequencing are likely enriched in functional binders. Higher frequency mutants are thus likely correlated with ACE2 function, and mutants that are less proteolytically sensitive are likely to be concentrated in this population.

We evaluated the correlation coefficient R between biological replicates “A” and “B” sorted by an average depth of coverage (**Fig 2B**). As expected, R decreases as the average depth of coverage of included mutations decreases. We do note that the correlation observed here (0.61-0.76 depending on the depth of coverage) is worse than typically observed in library sequencing experiments (>0.9) and limits our ability to make inferences about mutants that are more proteolytically susceptible than wild-type. Although we did not further investigate the reasons for lower correlation between replicates, one interpretation is the stringent maintenance of ACE binding in the reference population. Nevertheless, we found that at an average depth of coverage of 60 reads was sufficient to identify 40 mutants across 20 positions that had reproducible increases in their normalized enrichment ratios (**Fig 2B;** a heatmap of these positions are shown in **Fig 2C**). These hits are counted more frequently in the chymotrypsin-treated population than in the reference population, likely because they are resistant to chymotrypsin degradation.

The forty hits are largely distal to the RBM with the exception of multiple aliphatic mutations at N501 (N501F/W/M) (**Fig 2C, Fig 3A**). We neglected these hits as mutations at N501 are known to impact ACE2 affinity, including the N501Y mutation found in several Variants of Concern. The greatest surprise from the protease screen was that five of the eight mutations included in the RBD1 design (at positions 368, 410, 418, 438, 513) were found to have the WT residue at least somewhat enriched, suggesting that a reversion of that RBD1 mutation back to the wild-type sequence is stabilizing at that position (**Fig 2C, Fig 3A**). The overwhelming majority of other hits are concentrated in the underpacked core where the cryptic linoleic acid binding pocket occurs, most notably at positions 363 & 365 (**Fig 2C, Fig 3A**). There are a number of small to large aliphatic substitutions occurring at this pocket, including A363WPMILV, A397ML, I434F, and F392W. RBD1 contains a Y365W; both Y and W are disfavored relative to smaller aliphatics without hydrogen bond acceptors or donors. By contrast, the partially surface exposed V362 prefers the isosteric substitution threonine with hydrogen bonding potential.

**Figure 3:**
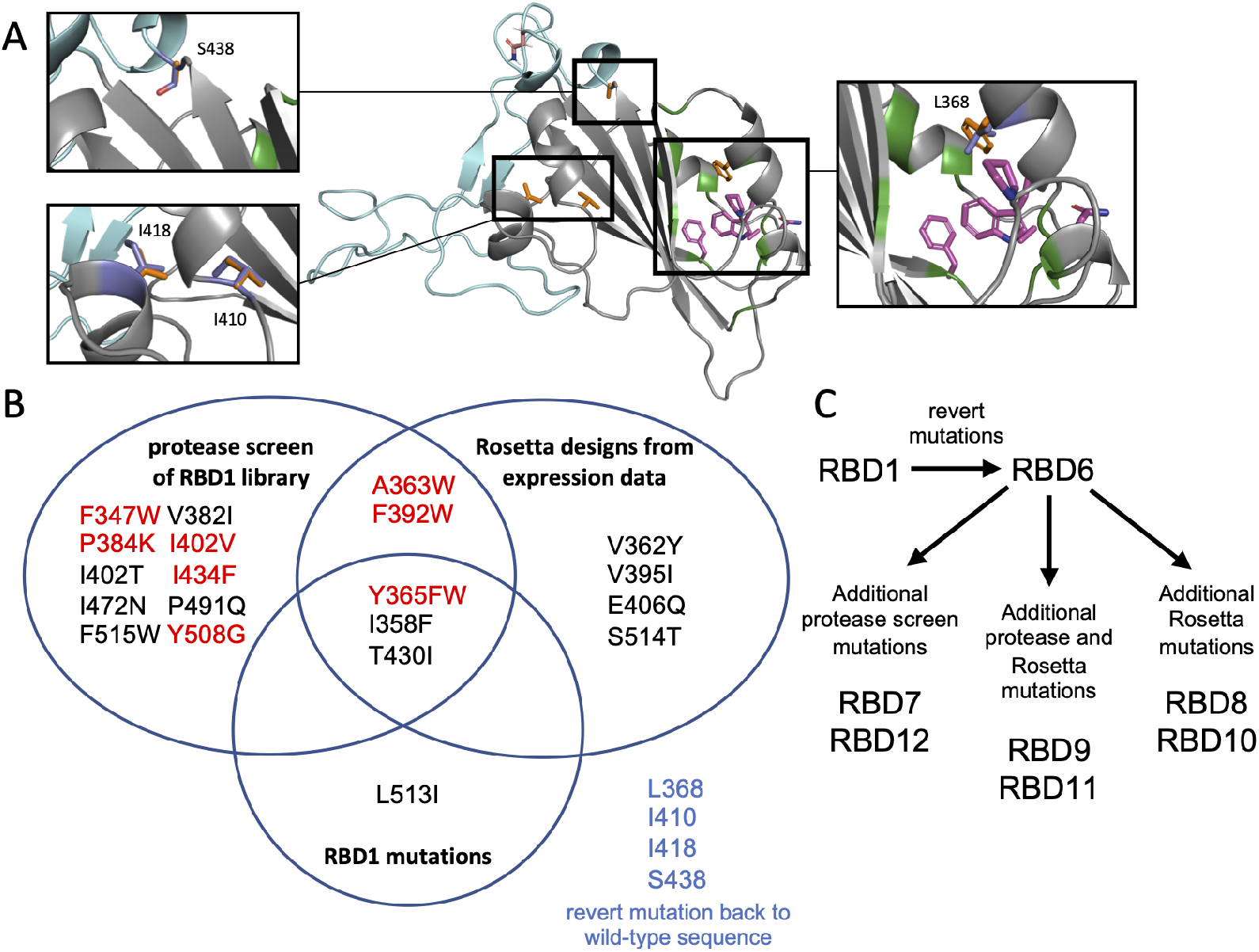
Sequence and structural identification of screening hits informs the next round of computational design. **A**. Selected stabilizing mutations identified by library protease screen were either reversions to wild-type of mutations incorporated into RBD1 (blue sticks; left inserts) or concentrated at or adjacent to the cryptic fatty acid binding pocket (pocket shown as green cartoon, and hits are shown as purple sticks; right insert). Orange sticks represent the original RBD1 residue. **B**. Distribution of identified stabilizing mutations incorporated into designs RBD6-RBD12. Hits with an average enrichment ratio > 1.0 and > 60 average read depth from library protease screen are colored red. **C**. Flowchart of process for distributing mutations among new designs.

We used the outputs from this deep mutational scan to inform the next set of computational designs (the full list of mutations used is shown as a Venn diagram in **Fig 3B**). The parsimonious design RBD6 is encoded by reversion of four mutations at positions 368, 410, 418, and 438. All other designs used this reversion as a base. We used Rosetta FastDesign on multiple RBD structures (see **Methods**) allowing either the RBD6 residue or a residue sourced from deep mutational scanning (**Fig 3C**). Two of the designs used all forty of the deep mutational scanning hits along with other potential hits just below the average depth of coverage. To add additional diversity of mutations, additional mutations were mined from a related deep mutational scanning dataset from Starr et al. (Starr et al. 2020) that quantified the effect of mutation on protein expression rather than stability. Since expression in yeast surface display is correlated with protein stability (Klesmith et al. 2017), core mutations that increase expression are likely to be stabilizing. Two designs incorporated mutations sourced from the Starr dataset. Finally, two designs considered mutations from both the Starr et al dataset along with the deep mutational scan from the current work.

All told, we created seven new designs containing a combination of nineteen different predicted stabilizing mutations (**Fig 4A**). Synthetic gblocks for the seven new designs were ordered and plasmids constructed as before. All designs displayed a c-myc epitope tag on the yeast surface, as determined by labeling with an anti-cmyc FITC secondary antibody. Two designs (RBD9, RBD12) bound neither ACE2 nor CR3022 at saturating concentrations, suggesting misfolding defects (**Fig 4A**). Two other designs (RBD7, RBD11) bound ACE2 but not CR3022, suggesting localized misfolding around the CR3022 epitope (**Fig 4A**). All four of these designs share the same sets of V382I and F515W mutations directly underneath the recognition surface of CR3022, suggesting that this combination of mutations likely results in local perturbation of the CR3022 epitope.

**Figure 4:**
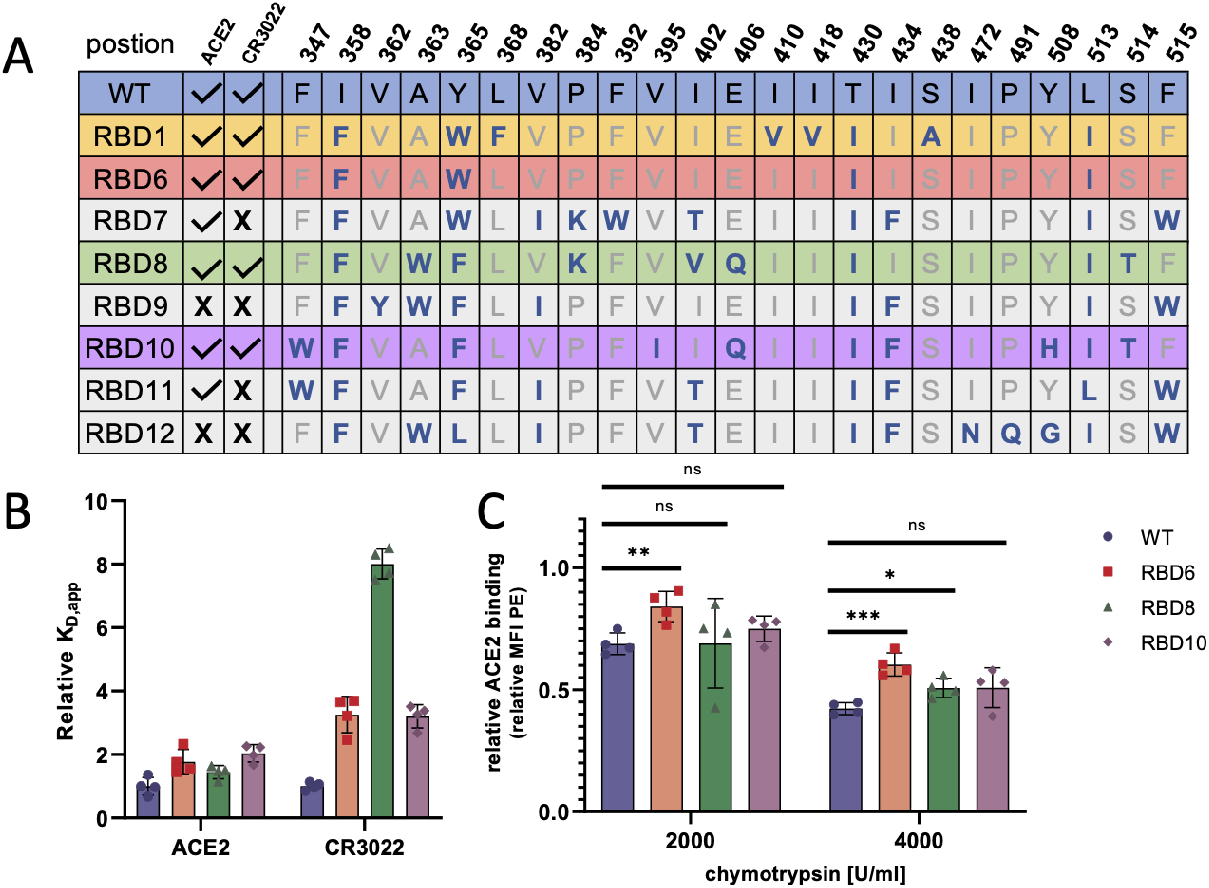
Core redesign produces a protein design with greater proteolytic stability compared to WT while maintaining recognition by binding proteins. **A**. The sequence diversity and binding assay results for the 2nd round of computational designs. Binding was assessed using cell surface labeling by ACE2-Fc (1 nM) and CR3022 (1 nM) of yeast displayed RBD designs. Only RBD6, RBD8, and RBD10 bound both ACE2 and CR3022. **B**. Relative apparent dissociation constants K_D, app_ for ACE2 and CR3022 for RBD6, RBD8, and RBD10 (n=4). K_D, app_ was assessed relative to WT RBD. **C**. ACE2 binding signal for WT, RBD6, RBD8, and RBD10 after treatment with increasing protease concentrations relative to no protease, measured by relative PE mean fluorescence intensity (RBD6 p-value = 7e-3 at 2000 U/ml, 5.3e-4 at 4000 U/ml chymotrypsin, RBD8 p-value = 0.011 at 4000 U/ml chymotrypsin, n=4).

Three of the seven new designs (RBD6, RBD8, RBD10) bound both ACE2 and CR3022. Cell surface titrations were performed as before, with all binding ACE2 comparable to WT and with slightly lower affinity for CR3022 (**Fig 4B**). Testing by the protease susceptibility assay shows that RBD6 (the RBD1 reversion) is significantly more stable than WT at both chymotrypsin concentrations tested (p-value = 7e-3 and 5.3e-4 at 2000 and 4000 U/ml chymotrypsin, respectively, n = 4) (**Fig 4C**). Design RBD8 was significantly more stable than WT only at the higher protease concentration of 4000 U/ml (p-value = 0.011, n = 4), while any differences in stability between design RBD10 and WT were not statistically significant at a significance threshold of 0.05 (**Fig 4C**). Based on this analysis we decided to express RBD6 and RBD8 recombinantly to determine whether in vitro properties correlated with the yeast surface display assays.

His-tagged WT, RBD6, and RBD8 constructs (S-RBD(333-537)) were expressed in *Pichia pastoris* and purified by affinity chromatography (Argentinian Covid Consortium et. al. 2020). All constructs purified as a single band at an apparent molecular weight of 31 kDa as visualized by SDS-PAGE, with Endo H treatment resulting in a single species migrating slightly faster at the predicted aglycosylated molecular weight of 28 kDa (**Fig S1A**). From this we conclude that all constructs are glycosylated at least at the lone N-linked glycan at position N343, consistent with previous reports of *Pichia* recombinant expression (Argentinian Covid Consortium et. al. 2020). For all constructs, we noticed the appearance of a slightly lower MW species visible by SDS-PAGE after extended storage (∼ 3-4 weeks) at 4°C (**Fig S1B**). We hypothesize that this additional species is a minor proteolysis product, as limited treatment with small amounts of thermolysin could reproduce the appearance of this band (**Fig S1C, S1D**).

We first assessed whether recombinant RBD variants could maintain binding to ACE2 and a panel of diverse antibodies that bind at distinct epitopes (CC12.7, CC12.1, CC6.29, CC6.32) (Rogers et al. 2020). Designs were screened for binding using an enzyme-linked immunosorbent assay (ELISA), in which recombinant RBD was immobilized on microtiter plates, incubated with varying concentrations of IgG or ACE-Fc, and then secondarily labeled with anti-human Fc-HRP conjugate. WT and designs assessed against ACE2-Fc results in a K_d,app_ between 1-2 nM (**Fig 5A-B**), and labeling with an IgG isotype control results in a K_d,app_ that could not be determined (>50 nM). RBD6 and RBD8 bound all antibodies on the panel with similar apparent affinities to WT (**Fig 5A-B**), suggesting strong structural conservation of the tertiary structure of the RBD protein domain.

**Figure 5:**
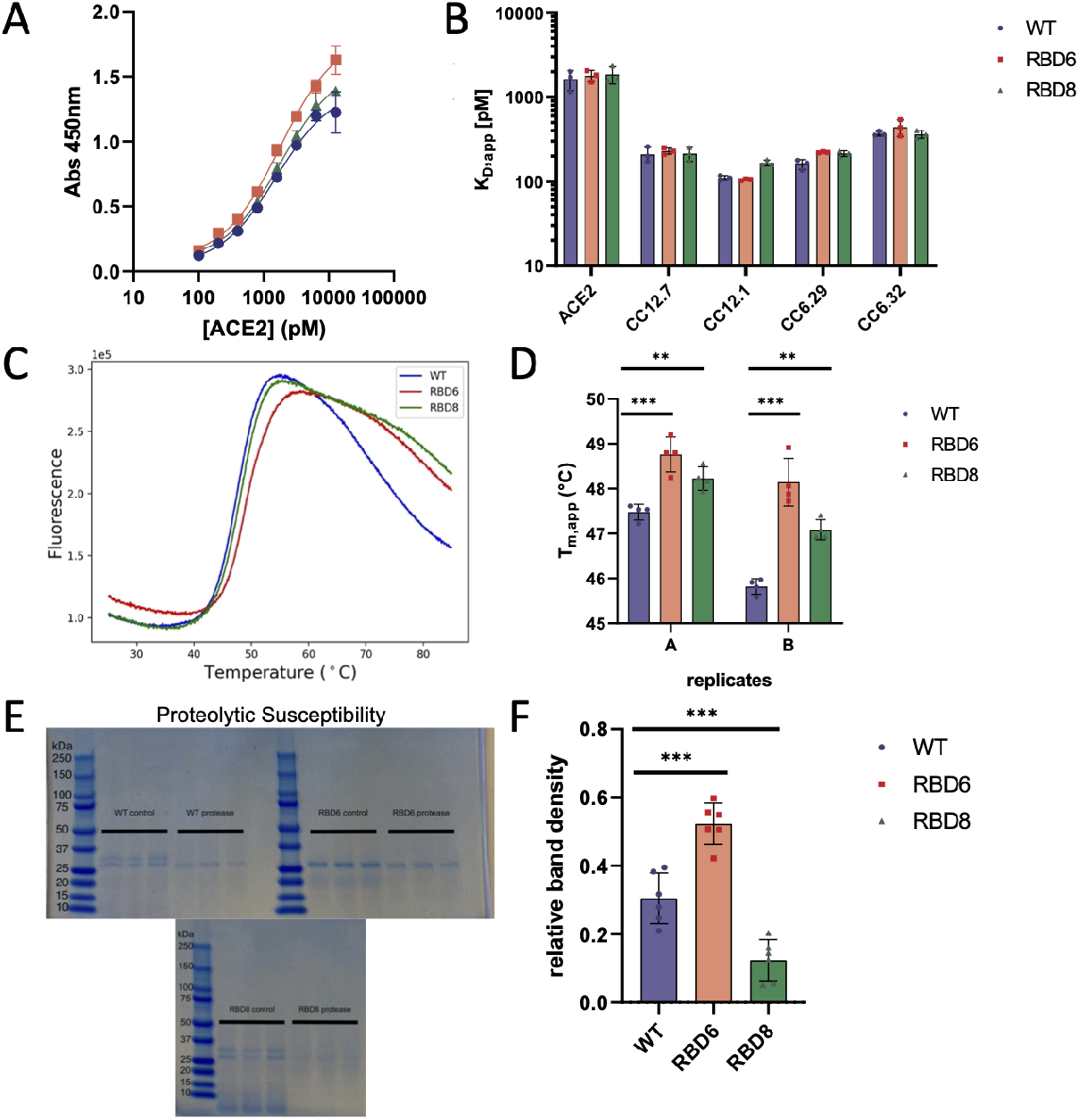
Recombinant RBD6 maintains binding to a wide range of neutralizing antibodies and has improved thermal and proteolytic stability compared to wild type RBD. **A**. hACE2-Fc titration curves for WT, RBD6, and RBD8 designs by ELISA (n=3). **B**. Calculated K_D,app_ for WT, RBD6, and RBD8 binding hACE2-Fc and antibodies CC12.7, CC12.1, CC6.29, and CC6.32 by ELISA (n=3). **C**. Fluorescence as a function of temperature using a SYPRO Orange thermal shift assay. **D**. Apparent melting temperature, T_m,app_, calculated from melting curves for WT, RBD6, and RBD8. Both designs show significantly higher Tm,app than WT for both replicates (RBD6: p-value = 1e-3, 5.7e-4; RBD8: p-value = 3.1e-3, 0.011) (n=4 each of 2 independent replicates). **E**. Representative SDS-PAGE gels showing WT, RBD6, and RBD8 protein after 1h incubation with the protease thermolysin. Thermolysin activity was quenched by addition of EDTA, and samples were then denatured and reduced. Gels were stained with SimplyBlue stain to visualize proteins. **F**. Quantification of protein band density after thermolysin relative to untreated control samples. RBD6 was significantly less sensitive to thermolysin treatment than WT (p-value = 0.00023), while RBD8 was more sensitive than WT to thermolysin treatment (p-value = 0.00090) (n=6).

We next assessed the apparent thermal stability (T_m,app_) of RBD6 and RBD8 compared to WT using a SYPRO-orange thermal shift assay (**Fig 5C**). Both RBD6 and RBD8 reported higher T_m,app_ than WT, at 1-2°C (p-value = 1e-3, 5.7e-4) and 0.5-1°C (p-value = 3.1e-3, 0.011), respectively (n = 4 in each replicate) (**Fig 5D**). Finally, we measured the relative proteolytic stability of the three constructs by incubating 1µg samples for 1 hour at 37°C with 193µM thermolysin. Denatured samples were visualized on an SDS-PAGE gel (**Fig 5E**). Quantification of the band intensity shows that more RBD6 protein remained after digestion with thermolysin than WT protein (p-value = 0.00023, n = 6), demonstrating that design RBD6 is more proteolytically stable than WT in addition to possessing increased thermal stability. Contrarily, design RBD8 was significantly less proteolytically stable than WT (p-value = 0.00090, n = 6), despite a modest increase in thermal stability (**Fig 5F**). From these sets of *in vitro* experiments we conclude that the design RBD6 maintains the tertiary surface topology of WT RBD, is more thermally stable, and is less proteolytically sensitive.

## Discussion

In this work we describe a case study on the thermal stabilization and proteolytic resistance of a SARS-CoV-2 S RBD variant engineered by computational design and deep mutational scanning. The RBD6 variant was still able to bind ACE2 and a panel of antibodies binding at distinct epitopes, showing that the increase in melting temperature and protease resistance did not appreciably change the exterior surface. As such, this is a stringent case of protein design and engineering distinct from other stabilization design and engineering strategies typically targeting surface or surface-proximal positions. Our design strategy is also distinct from the ‘S2P’ and ‘hexapro’ mutations on Spike (Pallesen et al. 2017, Hsieh et al. 2020) which stabilize certain protein conformations over others rather than necessarily stabilizing the folded state of the protein. Our approach could be applied to the computational stabilization of a wide range of proteins without requiring detailed knowledge of active sites or binding epitopes, particularly powerful for cases when there are multiple or unknown binding sites.

The four mutations encoded in the successful design RBD6 were I358F, Y365W, T430I, and I513L. While RBD6 resulted from a multi-point mutant reversion of four other nonoptimal residues from RBD1, our deep mutational scanning data indicate that many of these mutations may not be optimal. While tryptophan seems preferable to tyrosine at position 365, other aliphatic mutations like F without hydrogen bond donors or acceptors are superior to both. The mutation T430I removes a buried unsatisfied polar group. Our DMS data indicates that the isosteric aliphatic substitution of valine is slightly preferred over isoleucine at this position. Finally, both isoleucine and phenylalanine are slightly preferred over our originally designed I513L mutation in RBD1.

Similar to our work, the King group also found success mining the deep mutational scanning dataset of Starr et al (Starr et al. 2020, Ellis et al. 2021). Their choice to more narrowly focus on linoleic acid binding pocket mutations led to the creation of top designs containing only mutation F392W and mutations Y365F, F392W, and V395I, with melting temperatures of 1.9-2.4°C and 3.8-5.3°C above wild type, respectively (Ellis et. al. 2021). All of these mutations except V395I were identified at a high read depth by the protease library screen and show convergent strategies for RBD redesigns, even with distinct mutations. Thus we view our work as complementary to, and confirmatory of, the RBD redesign strategy employed by Ellis and colleagues.

This case study highlighted several areas in need of improvement for both computational and high-throughput protein engineering. In some respects, these issues are intertwined - higher throughput measurements combined with facile assembly of gene-length designs would allow for testing a richer diversity of design concepts than the two methods implemented here. On the computational side, the implementation of FuncLib led to a workable design whereas the three more aggressive FastDesign constructs did not display on the yeast surface. The number of mutations considered from the original sequence (8-18) is in the range of successful PROSS designs, as each mutation contributes perhaps 0.5-1°C increase in T_m_. In this specific case of the S RBD with only 80 or so mutable core residues, the number of mutations considered in the first round of designs was perhaps too aggressive and we would argue for the importance of parsimony in the set of initial future designs. The two-tiered screening strategy (design, experimentally characterize, then design again) resulted in a more stable RBD6 construct. However, RBD6 was a minimal reversion of the RBD1 construct, and more aggressive designs considering additional mutations ranged from minimally successful to failures. Two-tiered screening has a higher probability of success with a large diversity of the designs, such that potential failure modes are not shared between different designs.

On the experimental side, improving the hit identification from the protease assay used in the deep mutational scan would have improved the quality of the second set of designs. Our designs used both clear hits with average enrichment ratio greater than one, but also included some mutations with much lower average read depth (below 60) in order to increase diversity of designs. These borderline mutations generally were unsuccessful. Second, our scan considered only single point mutations from the RBD6 design. The majority of the hits were located at or adjacent to the linoleic acid binding pocket, and the experiment did not allow us to determine whether coupling of these individually stabilizing mutations were beneficial. Of course, introducing combinations of mutations increases the size of the library exponentially, and more clever library design would be needed to keep the number of variants to a manageable level. For this example, clear ways to limit library diversity would be to remove inviolable positions like the 8 cysteines involved in the 4 disulfide bonds, removal of positions like N501 where many residues are solvent exposed, and considering a reduced aliphatic alphabet (e.g. F,L,I,M,V,Y,A) rather than all 20 amino acids at completely buried positions. Such mutations could be efficiently programmed by existing combinatorial mutagenesis protocols (Kirby et al. 2021).

## Conclusion

In this work we used computational design and high throughput protein engineering to improve the thermal stability and reduce the proteolytic sensitivity of Wuhan Hu-1 S RBD without impacting its ability to recognize ACE2 and diverse antibodies targeting distinct epitopes.

## Supporting information

Complete Dataset for Manuscript Figures

## Supplementary Data

Supplementary data are available at *Protein Engineering, Design and Selection* online.

**Note S1**: Amino acid and nucleotide sequences of all RBD designs

**Figure S1**: Fluorescence-activated cell sorter gating for the deep mutational scanning screen

**Figure S2**: Degradation of purified RBD protein at 4°Cover time can be recapitulated by proteolytic degradation

**Table S1**: Plasmid List

**Table S2**: Primer List

**Table S3:** Library Statistics

## Funding

This work was supported by the National Science Foundation (CBET Award #2030221 to T.A.W.) and the National Institute Of Allergy And Infectious Diseases of the National Institutes of Health under Award Number R01AI141452 to T.A.W, the National Science Foundation Graduate Research Fellowship to A.C.L., and the NIH/CU Molecular Biophysics Program and NIH Biophysics Training Grant T32 GM-065103 to A.C.L. The content is solely the responsibility of the authors and does not necessarily represent the official views of the National Institutes of Health.

## Data Availability

Raw sequencing reads for this work have been deposited in the SRA (embargoed until publication). All processed deep sequencing runs and all raw data used for main text figures are available as a supplementary Microsoft Excel file (Leonard_complete_data.xlsx).

## Author Contributions

Designed proteins computationally: JW PJS ACL SJF TAW

Designed bench research: ACL TAW

Performed bench research: ACL AHE

Wrote manuscript: ACL TAW

## Conflicts of Interest

None noted.

## Notes

### Competing Interest Statement

The authors have declared no competing interest.

