## Supporting Information for "Stabilization of the SARS-CoV-2 Receptor Binding Domain by Protein Core Redesign and Deep Mutational Scanning"

3415 Colorado Avenue, Boulder, CO 80305

**Keywords**: SARS-CoV-2; Deep Mutational Scanning; Immunogen Design; FuncLib; Rosetta

**Note S1: RBD gene sequences**

>WT yeast surface display DNA sequence
ACAAACTTGTGCCCTTTTGGTGAAGTTTTTcaaGCCACCAGATTTGCATCTGTTTATGC
TTGGAACAGGAAGAGAATCAGCAACTGTGTTGCTGATTATTCTGTCCTATATAATTC
CGCATCATTTTCCACTTTTAAGTGTTATGGAGTGTCTCCTACTAAATTAAATGATCTC
TGCTTTACTAATGTCTATGCAGATTCATTTGTAATTAGAGGTGATGAAGTCAGACAA
ATCGCTCCAGGGCAAACTGGAAAGATTGCTGATTATAATTATAAATTACCAGATGAT
TTTACAGGCTGCGTTATAGCTTGGAATTCTAACAATCTTGATTCTAAGGTTGGTGGTA
ATTATAATTACCTGTATAGATTGTTTAGGAAGTCTAATCTCAAACCTTTTGAGAGAG
ATATTTCAACTGAAATCTATCAGGCCGGTAGCACACCTTGTAATGGTGTTGAAGGTT
TTAATTGTTACTTTCCTTTACAATCATATGGTTTCCAACCCACTAATGGTGTTGGTTA
CCAACCATACAGAGTAGTAGTACTTTCTTTTGAACTTCTACATGCACCAGCAACTGT
TTGTGGACCTAAAAAGTCTACTAATTTGGTTAAAAACAAA

>WT yeast surface display AA sequence
TNLCPFGEVFQATRFASVYAWNRKRISNCVADYSVLYNSASFSTFKCYGVSPTKLNDLC
FTNVYADSFVIRGDEVRQIAPGQTGKIADYNYKLPDDFTGCVIAWNSNNLDSKVGGNY
NYLYRLFRKSNLKPFERDISTEIYQAGSTPCNGVEGFNCYFPLQSYGFQPTNGVGYQPYR
VVVLSFELLHAPATVCGPKKSTNLVKNK

>RBD1 yeast surface display DNA sequence
accaacctgtgcccgtttggcgaagtgtttcaggcgacccgctttgcgagcgtgtatgcgtggaaccgcaaacgctttagcaactgcgtggc
ggattggagcgtgttttataacagcgcgagctttagcacctttaaatgctatggcgtgagcccgaccaaactgaacgatctgtgctttaccaac
gtgtatgcggatagctttgtgattcgcggcgatgaagtgcgccaggtggcgccgggccagaccggcaaagtggcggattataactataaa
ctgccggatgattttattggctgcgtgattgcgtggaacgcgaacaacctggatagcaaagtgggcggcaactataactatctgtatcgcctg
tttcgcaaaagcaacctgaaaccgtttgaacgcgatattagcaccgaaatttatcaggcgggcagcaccccgtgcaacggcgtggaaggct
ttaactgctattttccgctgcagagctatggctttcagccgaccaacggcgtgggctatcagccgtatcgcgtggtggtgattagctttgaact
gctgcatgcgccggcgaccgtgtgcggcccgaaaaaaagcaccaacctggtgaaaaacaaa

>RBD1 yeast surface display AA sequence
TNLCPFGEVFQATRFASVYAWNRKRFSNCVADWSVFYNSASFSTFKCYGVSPTKLNDL
CFTNVYADSFVIRGDEVRQVAPGQTGKVADYNYKLPDDFIGCVIAWNANNLDSKVGG
NYNYLYRLFRKSNLKPFERDISTEIYQAGSTPCNGVEGFNCYFPLQSYGFQPTNGVGYQP
YRVVVISFELLHAPATVCGPKKSTNLVKNK

>RBD2 yeast surface display DNA sequence
accaacctgtgcccgtttggcgaagtgtttcaggcgacccgctttgcgagcgtgtatgcgtggaaccgcaaacgcctgagcaactgcgtgg
cggattttagcgtgttttataacagcgcgagctttagcacctttaaatgctatggcgtgagcccgaccaaatttaacgatctgtgctttaccaacg
tgtatgcggatagctttgtgattcgcggcgatgaagtgcgccagattgcgccgggccagaccggcaaaattgcggattataactataaaattc
cggatgattttccgggctgcattgtggcgtggaacgcgaacaacctggatagcaaagtgggcggcaactataactatctgtatcgcctgtttc
gcaaaagcaacctgaaaccgtttgaacgcgatattagcaccgaaatttatcaggcgggcagcaccccgtgcaacggcgtggaaggcttta
actgctattttccgctgcagagctttggctttcagccgaccaacggcgtgggctatcagccgtatcgcgtggtggtgctgagctttgaactgct
gcatgcgccggcgaccgtgtgcggcccgaaaaaaagcaccaacctggtgaaaaacaaa

>RBD2 yeast surface display AA sequence
TNLCPFGEVFQATRFASVYAWNRKRLSNCVADFSVFYNSASFSTFKCYGVSPTKFNDLC
FTNVYADSFVIRGDEVRQIAPGQTGKIADYNYKIPDDFPGCIVAWNANNLDSKVGGNYN
YLYRLFRKSNLKPFERDISTEIYQAGSTPCNGVEGFNCYFPLQSFGFQPTNGVGYQPYRV
VVLSFELLHAPATVCGPKKSTNLVKNK

>RBD3 yeast surface display DNA sequence
accaacctgtgcccgtttggcgaactgtttcaggcgacccgctttgcggcggtgtatgcgtggaaccgcaaacgcctgagcaactgcgtgg
cggattatagcgtgctgtataacagcgcgagctttagcacctttaaatgctatggcgtgagcccgaccaaactgaacgatctgtgctgggtga
acgtgtatgtggatctgtttgtgattcgcggcgatgaagtgcgccagattgcgccgggccagaccggcaaaattgcggatctgaactataaa
ctgccggatgattttaccggctgcattattgtgtggaacgcgaacaacctggatgcgaaagtgggcggcaactataactatctgtatcgcctg
tttcgcaaaagcaacctgaaaccgctggaacgcgatattagcaccgaaatttatcaggcgggcagcaccccgtgcaacggcgtggaagg
ctttaactgctattttccgctgcagagctttggctggcagccgaccaacggcgtgggctatcagccgtttcgcgtggtggtgctgctgtttgaa
ctgctgcatgcgccggcgaccctgtgcggcccgaaaaaaagcaccaacctggtgaaaaacaaa

>RBD3 yeast surface display AA sequence
TNLCPFGELFQATRFAAVYAWNRKRLSNCVADYSVLYNSASFSTFKCYGVSPTKLNDL
CWVNVYVDLFVIRGDEVRQIAPGQTGKIADLNYKLPDDFTGCIIVWNANNLDAKVGGN
YNYLYRLFRKSNLKPLERDISTEIYQAGSTPCNGVEGFNCYFPLQSFGWQPTNGVGYQPF
RVVVLLFELLHAPATLCGPKKSTNLVKNK

>RBD4 yeast surface display DNA sequence
accaacctgtgcccgtttggcgaactgtttcaggcgacccgctttgcggcggtgtatgcgtggaaccgcaaacgcctgagcaactgcgtgg
cggattatagcgtgctgtataacagcgcgagctttagcacctttaaatgctatggcgtgagcccgaccaaactgaacgatctgtgctgggtga
acgtgtatattgatctgtttctgattcgcggcgatgaagtgcgccagattgcgccgggccagaccggcaaaattgcggatctgaactataaac
tgccggatgattttaccggctgcattattgtgtggaacgcgaacaacctggatgcgaaagtgggcggcaactataactatctgtatcgcctgtt
tcgcaaaagcaacctgaaaccgctggaacgcgatattagcaccgaaatttatcaggcgggcagcaccccgtgcaacggcgtggaaggct
ttaactgctattttccgctgcagagctttggctttcagccgaccaacggcgtgggctatcagccgtttcgcgtggtggtgctgctgtttgaactg
ctgcatgcgccggcgaccctgtgcggcccgaaaaaaagcaccaacctggtgaaaaacaaa

>RBD4 yeast surface display AA sequence
TNLCPFGELFQATRFAAVYAWNRKRLSNCVADYSVLYNSASFSTFKCYGVSPTKLNDL
CWVNVYIDLFLIRGDEVRQIAPGQTGKIADLNYKLPDDFTGCIIVWNANNLDAKVGGNY
NYLYRLFRKSNLKPLERDISTEIYQAGSTPCNGVEGFNCYFPLQSFGFQPTNGVGYQPFR
VVVLLFELLHAPATLCGPKKSTNLVKNK

>RBD5 yeast surface display DNA sequence
accaacctgtgcccgtttggcgaactgtttcaggcgacccgctttgcggcggtgtatgcgtggaaccgcaaacgctttagcaactgcgtggc
ggattatagcgtgctgtataacagcgcgagctttagcacctttaaatgctatggcgtgagcccgaccaaactgaacgatctgtgctttgtgaac
gtgtatgtggatctgtttctgattcgcggcgatgaagtgcgccagattgcgccgggccagaccggcaaaattgcggatctgaactataaact
gccggatgattggaccggctgcattattgtgtggaacgcgaacaacctggatgcgaaagtgggcggcaactataactatctgtatcgcctgt
ttcgcaaaagcaacctgaaaccgctggaacgcgatattagcaccgaaatttatcaggcgggcagcaccccgtgcaacggcgtggaaggc
tttaactgctattttccgctgcagagctttggctttcagccgaccaacggcgtgggctatctgccgtttcgcgtggtggtgctgatttttgaactg
ctgcatgcgccggcgaccatttgcggcccgaaaaaaagcaccaacctggtgaaaaacaaa

>RBD5 yeast surface display AA sequence
TNLCPFGELFQATRFAAVYAWNRKRFSNCVADYSVLYNSASFSTFKCYGVSPTKLNDL
CFVNVYVDLFLIRGDEVRQIAPGQTGKIADLNYKLPDDWTGCIIVWNANNLDAKVGGN
YNYLYRLFRKSNLKPLERDISTEIYQAGSTPCNGVEGFNCYFPLQSFGFQPTNGVGYLPF
RVVVLIFELLHAPATICGPKKSTNLVKNK

>RBD6 yeast surface display DNA sequence
accaacctgtgcccgtttggcgaagtgtttcaggcgacccgctttgcgagcgtgtatgcgtggaaccgcaaacgctttagcaactgcgtggc
ggattggagcgtgctgtataacagcgcgagctttagcacctttaaatgctatggcgtgagcccgaccaaactgaacgatctgtgctttaccaa
cgtgtatgcggatagctttgtgattcgcggcgatgaagtgcgccagattgcgccgggccagaccggcaaaattgcggattataactataaac
tgccggatgattttattggctgcgtgattgcgtggaacagcaacaacctggatagcaaagtgggcggcaactataactatctgtatcgcctgtt
tcgcaaaagcaacctgaaaccgtttgaacgcgatattagcaccgaaatttatcaggcgggcagcaccccgtgcaacggcgtggaaggcttt
aactgctattttccgctgcagagctatggctttcagccgaccaacggcgtgggctatcagccgtatcgcgtggtggtgattagctttgaactgc
tgcatgcgccggcgaccgtgtgcggcccgaaaaaaagcaccaacctggtgaaaaacaaa

>RBD6 yeast surface display AA sequence
TNLCPFGEVFQATRFASVYAWNRKRFSNCVADWSVLYNSASFSTFKCYGVSPTKLNDL
CFTNVYADSFVIRGDEVRQIAPGQTGKIADYNYKLPDDFIGCVIAWNSNNLDSKVGGNY
NYLYRLFRKSNLKPFERDISTEIYQAGSTPCNGVEGFNCYFPLQSYGFQPTNGVGYQPYR
VVVISFELLHAPATVCGPKKSTNLVKNK

>RBD7 yeast surface display DNA sequence
accaacctgtgcccgtttggcgaagtgtttcaggcgacccgctttgcgagcgtgtatgcgtggaaccgcaaacgctttagcaactgcgtggc
ggattggagcgtgctgtataacagcgcgagctttagcacctttaaatgctatggcattagcaaaaccaaactgaacgatctgtgctggaccaa
cgtgtatgcggatagctttgtgacccgcggcgatgaagtgcgccagattgcgccgggccagaccggcaaaattgcggattataactataaa
ctgccggatgattttattggctgcgtgtttgcgtggaacagcaacaacctggatagcaaagtgggcggcaactataactatctgtatcgcctgt
ttcgcaaaagcaacctgaaaccgtttgaacgcgatattagcaccgaaatttatcaggcgggcagcaccccgtgcaacggcgtggaaggctt
taactgctattttccgctgcagagctatggctttcagccgaccaacggcgtgggctatcagccgtatcgcgtggtggtgattagctgggaact
gctgcatgcgccggcgaccgtgtgcggcccgaaaaaaagcaccaacctggtgaaaaacaaa

>RBD7 yeast surface display AA sequence
TNLCPFGEVFQATRFASVYAWNRKRFSNCVADWSVLYNSASFSTFKCYGISKTKLNDLC
WTNVYADSFVTRGDEVRQIAPGQTGKIADYNYKLPDDFIGCVFAWNSNNLDSKVGGNY
NYLYRLFRKSNLKPFERDISTEIYQAGSTPCNGVEGFNCYFPLQSYGFQPTNGVGYQPYR
VVVISWELLHAPATVCGPKKSTNLVKNK

>RBD8 yeast surface display DNA sequence
accaacctgtgcccgtttggcgaagtgtttcaggcgacccgctttgcgagcgtgtatgcgtggaaccgcaaacgctttagcaactgcgtgtg
ggattttagcgtgctgtataacagcgcgagctttagcacctttaaatgctatggcgtgagcaaaaccaaactgaacgatctgtgctttaccaac
gtgtatgcggatagctttgtggtgcgcggcgatcaggtgcgccagattgcgccgggccagaccggcaaaattgcggattataactataaac
tgccggatgattttattggctgcgtgattgcgtggaacagcaacaacctggatagcaaagtgggcggcaactataactatctgtatcgcctgtt
tcgcaaaagcaacctgaaaccgtttgaacgcgatattagcaccgaaatttatcaggcgggcagcaccccgtgcaacggcgtggaaggcttt
aactgctattttccgctgcagagctatggctttcagccgaccaacggcgtgggctatcagccgtatcgcgtggtggtgattacctttgaactgc
tgcatgcgccggcgaccgtgtgcggcccgaaaaaaagcaccaacctggtgaaaaacaaa

>RBD8 yeast surface display AA sequence
TNLCPFGEVFQATRFASVYAWNRKRFSNCVWDFSVLYNSASFSTFKCYGVSKTKLNDL
CFTNVYADSFVVRGDQVRQIAPGQTGKIADYNYKLPDDFIGCVIAWNSNNLDSKVGGN
YNYLYRLFRKSNLKPFERDISTEIYQAGSTPCNGVEGFNCYFPLQSYGFQPTNGVGYQPY
RVVVITFELLHAPATVCGPKKSTNLVKNK

>RBD9 yeast surface display DNA sequence
accaacctgtgcccgtttggcgaagtgtttcaggcgacccgctttgcgagcgtgtatgcgtggaaccgcaaacgctttagcaactgctattgg
gattttagcgtgctgtataacagcgcgagctttagcacctttaaatgctatggcattagcccgaccaaactgaacgatctgtgctttaccaacgt
gtatgcggatagctttgtgattcgcggcgatgaagtgcgccagattgcgccgggccagaccggcaaaattgcggattataactataaactgc
cggatgattttattggctgcgtgtttgcgtggaacagcaacaacctggatagcaaagtgggcggcaactataactatctgtatcgcctgtttcg
caaaagcaacctgaaaccgtttgaacgcgatattagcaccgaaatttatcaggcgggcagcaccccgtgcaacggcgtggaaggctttaa
ctgctattttccgctgcagagctatggctttcagccgaccaacggcgtgggctatcagccgtatcgcgtggtggtgattagctgggaactgct
gcatgcgccggcgaccgtgtgcggcccgaaaaaaagcaccaacctggtgaaaaacaaa

>RBD9 yeast surface display AA sequence
TNLCPFGEVFQATRFASVYAWNRKRFSNCYWDFSVLYNSASFSTFKCYGISPTKLNDLC
FTNVYADSFVIRGDEVRQIAPGQTGKIADYNYKLPDDFIGCVFAWNSNNLDSKVGGNY
NYLYRLFRKSNLKPFERDISTEIYQAGSTPCNGVEGFNCYFPLQSYGFQPTNGVGYQPYR
VVVISWELLHAPATVCGPKKSTNLVKNK

>RBD10 yeast surface display DNA sequence
accaacctgtgcccgtttggcgaagtgtttcaggcgacccgctgggcgagcgtgtatgcgtggaaccgcaaacgctttagcaactgcgtgg
cggattttagcgtgctgtataacagcgcgagctttagcacctttaaatgctatggcgtgagcccgaccaaactgaacgatctgtgctttaccaa
catttatgcggatagctttgtgattcgcggcgatcaggtgcgccagattgcgccgggccagaccggcaaaattgcggattataactataaact
gccggatgattttattggctgcgtgtttgcgtggaacagcaacaacctggatagcaaagtgggcggcaactataactatctgtatcgcctgttt
cgcaaaagcaacctgaaaccgtttgaacgcgatattagcaccgaaatttatcaggcgggcagcaccccgtgcaacggcgtggaaggcttt
aactgctattttccgctgcagagctatggctttcagccgaccaacggcgtgggctatcagccgcatcgcgtggtggtgattacctttgaactg
ctgcatgcgccggcgaccgtgtgcggcccgaaaaaaagcaccaacctggtgaaaaacaaa

>RBD10 yeast surface display AA sequence
TNLCPFGEVFQATRWASVYAWNRKRFSNCVADFSVLYNSASFSTFKCYGVSPTKLNDL
CFTNIYADSFVIRGDQVRQIAPGQTGKIADYNYKLPDDFIGCVFAWNSNNLDSKVGGNY
NYLYRLFRKSNLKPFERDISTEIYQAGSTPCNGVEGFNCYFPLQSYGFQPTNGVGYQPHR
VVVITFELLHAPATVCGPKKSTNLVKNK

>RBD11 yeast surface display DNA sequence
accaacctgtgcccgtttggcgaagtgtttcaggcgacccgctgggcgagcgtgtatgcgtggaaccgcaaacgctttagcaactgcgtgg
cggattttagcgtgctgtataacagcgcgagctttagcacctttaaatgctatggcattagcccgaccaaactgaacgatctgtgctttaccaac
gtgtatgcggatagctttgtgacccgcggcgatgaagtgcgccagattgcgccgggccagaccggcaaaattgcggattataactataaac
tgccggatgattttattggctgcgtgtttgcgtggaacagcaacaacctggatagcaaagtgggcggcaactataactatctgtatcgcctgttt
cgcaaaagcaacctgaaaccgtttgaacgcgatattagcaccgaaatttatcaggcgggcagcaccccgtgcaacggcgtggaaggcttt
aactgctattttccgctgcagagctatggctttcagccgaccaacggcgtgggctatcagccgtatcgcgtggtggtgctgagctgggaact
gctgcatgcgccggcgaccgtgtgcggcccgaaaaaaagcaccaacctggtgaaaaacaaa

>RBD11 yeast surface display AA sequence
TNLCPFGEVFQATRWASVYAWNRKRFSNCVADFSVLYNSASFSTFKCYGISPTKLNDLC
FTNVYADSFVTRGDEVRQIAPGQTGKIADYNYKLPDDFIGCVFAWNSNNLDSKVGGNY
NYLYRLFRKSNLKPFERDISTEIYQAGSTPCNGVEGFNCYFPLQSYGFQPTNGVGYQPYR
VVVLSWELLHAPATVCGPKKSTNLVKNK

>RBD12 yeast surface display DNA sequence
accaacctgtgcccgtttggcgaagtgtttcaggcgacccgctttgcgagcgtgtatgcgtggaaccgcaaacgctttagcaactgcgtgtg
ggatctgagcgtgctgtataacagcgcgagctttagcacctttaaatgctatggcattagcccgaccaaactgaacgatctgtgctttaccaac
gtgtatgcggatagctttgtgacccgcggcgatgaagtgcgccagattgcgccgggccagaccggcaaaattgcggattataactataaac
tgccggatgattttattggctgcgtgtttgcgtggaacagcaacaacctggatagcaaagtgggcggcaactataactatctgtatcgcctgttt
cgcaaaagcaacctgaaaccgtttgaacgcgatattagcaccgaaaactatcaggcgggcagcaccccgtgcaacggcgtggaaggcttt
aactgctattttcagctgcagagctatggctttcagccgaccaacggcgtgggctatcagccgggccgcgtggtggtgattagctgggaact
gctgcatgcgccggcgaccgtgtgcggcccgaaaaaaagcaccaacctggtgaaaaacaaa

>RBD12 yeast surface display AA sequence
TNLCPFGEVFQATRFASVYAWNRKRFSNCVWDLSVLYNSASFSTFKCYGISPTKLNDLC
FTNVYADSFVTRGDEVRQIAPGQTGKIADYNYKLPDDFIGCVFAWNSNNLDSKVGGNY
NYLYRLFRKSNLKPFERDISTENYQAGSTPCNGVEGFNCYFQLQSYGFQPTNGVGYQPG
RVVVISWELLHAPATVCGPKKSTNLVKNK

>WT Pichia pastoris expression DNA sequence
ACCAATCTTTGTCCATTTGGAGAAGTTTTCAACGCAACGAGATTCGCAAGTGTCTAC
GCCTGGAACCGTAAGAGGATAAGTAATTGTGTGGCTGACTATTCAGTGTTGTATAAC
TCTGCCTCATTTTCCACCTTCAAATGCTACGGCGTATCTCCAACAAAGCTAAACGATT
TGTGTTTCACCAATGTATACGCCGATTCCTTTGTCATAAGGGGAGATGAGGTACGTC
AAATAGCCCCAGGCCAGACGGGAAAAATAGCTGATTATAACTATAAATTGCCTGAT
GACTTTACAGGCTGCGTTATCGCATGGAACTCAAACAATTTGGACTCTAAGGTGGGC
GGCAATTACAATTATCTTTACAGACTTTTCAGGAAATCCAATCTAAAGCCATTTGAG
CGTGATATTTCTACAGAAATATATCAAGCTGGTTCTACCCCATGTAATGGAGTAGAG
GGTTTTAATTGTTATTTCCCATTGCAATCCTATGGTTTTCAACCTACTAATGGAGTGG
GATACCAGCCTTATAGGGTCGTGGTGCTTTCTTTTGAATTACTTCATGCCCCCGCTAC
CGTTTGCGGTCCCAAAAAATCTACAAACCTAGTTAAGAACAAA

>WT Pichia pastoris expression AA sequence
TNLCPFGEVFNATRFASVYAWNRKRISNCVADYSVLYNSASFSTFKCYGVSPTKLNDLC
FTNVYADSFVIRGDEVRQIAPGQTGKIADYNYKLPDDFTGCVIAWNSNNLDSKVGGNY
NYLYRLFRKSNLKPFERDISTEIYQAGSTPCNGVEGFNCYFPLQSYGFQPTNGVGYQPYR
VVVLSFELLHAPATVCGPKKSTNLVKNK

>RBD6 Pichia pastoris expression DNA sequence
ACCAATCTTTGTCCATTTGGAGAAGTTTTCAACGCAACGAGATTCGCAAGTGTCTAC
GCCTGGAACCGTAAGAGGTTCAGTAATTGTGTGGCTGACTGGTCAGTGTTGTATAAC
TCTGCCTCATTTTCCACCTTCAAATGCTACGGCGTATCTCCAACAAAGCTAAACGATT
TGTGTTTCACCAATGTATACGCCGATTCCTTTGTCATAAGGGGAGATGAGGTACGTC
AAATAGCCCCAGGCCAGACGGGAAAAATAGCTGATTATAACTATAAATTGCCTGAT

GACTTTATAGGCTGCGTTATCGCATGGAACTCAAACAATTTGGACTCTAAGGTGGGC
GGCAATTACAATTATCTTTACAGACTTTTCAGGAAATCCAATCTAAAGCCATTTGAG
CGTGATATTTCTACAGAAATATATCAAGCTGGTTCTACCCCATGTAATGGAGTAGAG
GGTTTTAATTGTTATTTCCCATTGCAATCCTATGGTTTTCAACCTACTAATGGAGTGG
GATACCAGCCTTATAGGGTCGTGGTGATATCTTTTGAATTACTTCATGCCCCCGCTAC
CGTTTGCGGTCCCAAAAAATCTACAAACCTAGTTAAGAACAAA

>RBD6 Pichia pastoris expression AA sequence
TNLCPFGEVFNATRFASVYAWNRKRFSNCVADWSVLYNSASFSTFKCYGVSPTKLNDL
CFTNVYADSFVIRGDEVRQIAPGQTGKIADYNYKLPDDFIGCVIAWNSNNLDSKVGGNY
NYLYRLFRKSNLKPFERDISTEIYQAGSTPCNGVEGFNCYFPLQSYGFQPTNGVGYQPYR
VVVISFELLHAPATVCGPKKSTNLVKNK

>RBD8 Pichia pastoris expression DNA sequence
ACCAATCTTTGTCCATTTGGAGAAGTTTTCAACGCAACGAGATTCGCAA
GTGTCTACGCCTGGAACCGTAAGAGGTTCAGTAATTGTGTGTGGGACTT
CTCAGTGTTGTATAACTCTGCCTCATTTTCCACCTTCAAATGCTACGGCG
TATCTAAAACAAAGCTAAACGATTTGTGTTTCACCAATGTATACGCCGA
TTCCTTTGTCGTAAGGGGAGATCAGGTACGTCAAATAGCCCCAGGCCA
GACGGGAAAAATAGCTGATTATAACTATAAATTGCCTGATGACTTTATA
GGCTGCGTTATCGCATGGAACTCAAACAATTTGGACTCTAAGGTGGGC
GGCAATTACAATTATCTTTACAGACTTTTCAGGAAATCCAATCTAAAGC
CATTTGAGCGTGATATTTCTACAGAAATATATCAAGCTGGTTCTACCCC
ATGTAATGGAGTAGAGGGTTTTAATTGTTATTTCCCATTGCAATCCTAT
GGTTTTCAACCTACTAATGGAGTGGGATACCAGCCTTATAGGGTCGTGG
TGATAACCTTTGAATTACTTCATGCCCCCGCTACCGTTTGCGGTCCCAA
AAAATCTACAAACCTAGTTAAGAACAAA

>RBD8 Pichia pastoris expression AA sequence
TNLCPFGEVFNATRFASVYAWNRKRFSNCVWDFSVLYNSASFSTFKCYGV
SKTKLNDLCFTNVYADSFVVRGDQVRQIAPGQTGKIADYNYKLPDDFIGC
VIAWNSNNLDSKVGGNYNYLYRLFRKSNLKPFERDISTEIYQAGSTPCNGV
EGFNCYFPLQSYGFQPTNGVGYQPYRVVVITFELLHAPATVCGPKKSTNLV
KNK

#### Table S1: Plasmid List

| Name | Description | *E. coli* marker | *S. cerevisiae* marker | *P. pastoris* marker | Source |
|---|---|---|---|---|---|
| Production_pETCON_V4_B1A2 (pETconV4) | Yeast surface display plasmid optimized for protease susceptibility assay | Amp | TRP1 | | Maguire et. al., 2021 |
| pJS699 | S-RBD(333-537)-N343Q | | | | Banach et al., 2021 |
| pACL002 | Design 00481 in pETconV4 plasmid | Amp | TRP1 | | In house |
| pACL003 | Design 01063 in pETconV4 plasmid | Amp | TRP1 | | In house |
| pACL004 | Design 05626 in pETconV4 plasmid | Amp | TRP1 | | In house |
| pACL005 | Design F1 in pETconV4 plasmid | Amp | TRP1 | | In house |
| pACL006 | Design F2 in pETconV4 | Amp | TRP1 | | In house |
| pACL007 | WT RBD sequence in pETconV4 | Amp | TRP1 | | In house |
| pACL008 | pACL005 with BbvCI restriction sites | Amp | TRP1 | | In house |
| pACL009 | Design 1 in pETconV4 plasmid | Amp | TRP1 | | In house |
| pACL010 | Design 2 in pETconV4 plasmid | Amp | TRP1 | | In house |
| pACL011 | Design 3 in pETconV4 plasmid | Amp | TRP1 | | In house |
| pACL012 | Design 4 in pETconV4 plasmid | Amp | TRP1 | | In house |
| pACL013 | Design 5 in pETconV4 plasmid | Amp | TRP1 | | In house |
| pACL014 | Design 6 in pETconV4 plasmid | Amp | TRP1 | | In house |
| pACL015 | Design 7 in pETconV4 plasmid | Amp | TRP1 | | In house |
| pPICZ$\alpha$A | Plasmid for secreted expression of recombinant proteins in *P. pastoris*, reference frame A | Zeocin | | Zeocin | ThermoFisher Cat#V19520 |
| pACL020 | Wild type in pPICZ$\alpha$A plasmid | Zeocin | | Zeocin | In house |
| pACL021 | Design 1 in pPICZ$\alpha$A plasmid | Zeocin | | Zeocin | In house |
| pACL022 | Design 3 in pPICZ$\alpha$A plasmid | Zeocin | | Zeocin | In house |
| pACL023 | Design 5 in pPICZ$\alpha$A plasmid | Zeocin | | Zeocin | In house |

#### Table S2: Primer List

| Name | Sequence 5'->3' | Description |
|---|---|---|
| forward_pJS699_RBD_pETconV4 | gagggtcggcttcgcatatgacaaacttatgcccttttgg | Add pETconV4 homologous region to pJS699 |
| reverse_pJS699_RBD_pETconV4-rev | ycggaacctccaccctcgagtttgtttttaaccaaattagtagactt | |
| pETconV4_seq_rev | gtgggaacaaagtcgattttgTTACATC | Primer for Sanger sequencing RBD insert in pETconV4-based plasmids |
| pETconV4_BbvCI_sequencing | gcttcccggcaacaattaatagactg | Primer for Sanger sequencing to confirm insertion of BbvCI site in pETconV4-based plasmids |
| add_BbvCI_forw_pETconV4 | gcaatcagaccaagtttactcatatatactttag | Insert BbvCI cut site using Q5 site directed mutagenesis |
| add_BbvCI_rev_pETconV4 | tgaggcagttaccaatgcttaatcagtgag | |
| RBD-F1_tile1_F_Illumina | gttcagagttctacagtccgacgatcCGGAGGGTCGGCTTCGCATATG | Tile 1 forward inner primer with adapter for Illumina sequencing |
| RBD-F1_tile1_R_Illumina | ccttggcacccgagaattccacgcgttccacgcaatcacgc | Tile 1 reverse inner primer with adapter for Illumina sequencing |
| RBD-F1_tile2_F_Illumina | gttcagagttctacagtccgacgatctaaactgccggatgattttattggc | Tile 2 forward inner primer with adapter for Illumina sequencing |
| RBD-F1_tile2_R_Illumina | ccttggcacccgagaattccacaggttggtgcttttttcgg | Tile 2 reverse inner primer with adapter for Illumina sequencing |
| colonyPCR_pichiaRBD_rev | CTACTCCATTACATGGGGTAG | Colony PCR primer for pPICZ$\alpha$-based plasmids |
| pPICZalpha_seq_rev | GAACTGAGGAACAGTCATGTC | Primer for Sanger sequencing RBD insert in pPICZ$\alpha$ -based plasmids |
| Q343N_SDM_WT_D1_D3_F | AACGCAACGAGATTCGCAAGTGTC | Site-directed mutagenesis primers to revert N343Q mutation |
| Q343N_SDM_R | GAAAACTTCTCCAAATGGACAAAGATTG | |

| | | |
|---|---|---|
| RBD_F1-C4-NNK | CATATGACCAACCTGNNKCCGTTTGGCGAAGTG | Degenerate primer set for nicking mutagenesis |
| RBD_F1-F6-NNK | ACCAACCTGTGCCCGNNKGGCGAAGTGTTTCAG | |
| RBD_F1-V9-NNK | TGCCCGTTTGGCGAANNKTTTCAGGCGACCCGC | |
| RBD_F1-F10-NNK | CCGTTTGGCGAAGTGNNKCAGGCGACCCGCTTT | |
| RBD_F1-F15-NNK | TTTCAGGCGACCCGCNNKGCGAGCGTGTATGCG | |
| RBD_F1-S17-NNK | GCGACCCGCTTTGCGNNKGTGTATGCGTGGAAC | |

| | |
|---|---|
| RBD_F1-V18-NNK | ACCCGCTTTGCGAGCNNKTATGCGTGGAACCGC |
| RBD_F1-Y19-NNK | CGCTTTGCGAGCGTGNNKGCGTGGAACCGCAAA |
| RBD_F1-A20-NNK | TTTGCGAGCGTGTATNNKTGGAACCGCAAACGC |
| RBD_F1-W21-NNK | GCGAGCGTGTATGCGNNKAACCGCAAACGCTTT |
| RBD_F1-F26-NNK | TGGAACCGCAAACGCNNKAGCAACTGCGTGGCG |
| RBD_F1-C29-NNK | AAACGCTTTAGCAACNNKGTGGCGGATTGGAGC |
| RBD_F1-V30-NNK | CGCTTTAGCAACTGCNNKGCGGATTGGAGCGTG |
| RBD_F1-A31-NNK | TTTAGCAACTGCGTGNNKGATTGGAGCGTGTTT |
| RBD_F1-W33-NNK | AACTGCGTGGCGGATNNKAGCGTGTTTTATAAC |
| RBD_F1-F36-NNK | GCGGATTGGAGCGTGNNKTATAACAGCGCGAGC |
| RBD_F1-F42-NNK | TATAACAGCGCGAGCNNKAGCACCTTTAAATGC |
| RBD_F1-F45-NNK | GCGAGCTTTAGCACCNNKAAATGCTATGGCGTG |
| RBD_F1-C47-NNK | TTTAGCACCTTTAAANNKTATGGCGTGAGCCCG |
| RBD_F1-V50-NNK | TTTAAATGCTATGGCNNKAGCCCGACCAAACTG |
| RBD_F1-P52-NNK | TGCTATGGCGTGAGCNNKACCAAACTGAACGAT |
| RBD_F1-L55-NNK | GTGAGCCCGACCAAANNKAACGATCTGTGCTTT |
| RBD_F1-N56-NNK | AGCCCGACCAAACTGNNKGATCTGTGCTTTACC |
| RBD_F1-F60-NNK | CTGAACGATCTGTGCNNKACCAACGTGTATGCG |
| RBD_F1-T61-NNK | AACGATCTGTGCTTTNNKAACGTGTATGCGGAT |
| RBD_F1-N62-NNK | GATCTGTGCTTTACCNNKGTGTATGCGGATAGC |
| RBD_F1-V63-NNK | CTGTGCTTTACCAACNNKTATGCGGATAGCTTT |
| RBD_F1-Y64-NNK | TGCTTTACCAACGTGNNKGCGGATAGCTTTGTG |
| RBD_F1-A65-NNK | TTTACCAACGTGTATNNKGATAGCTTTGTGATT |
| RBD_F1-D66-NNK | ACCAACGTGTATGCGNNKAGCTTTGTGATTCGC |
| RBD_F1-S67-NNK | AACGTGTATGCGGATNNKTTTGTGATTCGCGGC |
| RBD_F1-F68-NNK | GTGTATGCGGATAGCNNKGTGATTCGCGGCGAT |
| RBD_F1-V69-NNK | TATGCGGATAGCTTTNNKATTCGCGGCGATGAA |
| RBD_F1-I70-NNK | GCGGATAGCTTTGTGNNKCGCGGCGATGAAGTG |
| RBD_F1-G72-NNK | AGCTTTGTGATTCGCNNKGATGAAGTGCGCCAG |
| RBD_F1-E74-NNK | GTGATTCGCGGCGATNNKGTGCGCCAGGTGGCG |
| RBD_F1-V75-NNK | ATTCGCGGCGATGAANNKCGCCAGGTGGCGCCG |
| RBD_F1-Q77-NNK | GGCGATGAAGTGCGCNNKGTGGCGCCGGGCCAG |
| RBD_F1-V78-NNK | GATGAAGTGCGCCAGNNKGCGCCGGGCCAGACC |
| RBD_F1-A79-NNK | GAAGTGCGCCAGGTGNNKCCGGGCCAGACCGGC |
| RBD_F1-P80-NNK | GTGCGCCAGGTGGCGNNKGGCCAGACCGGCAAA |
| RBD_F1-V86-NNK | GGCCAGACCGGCAAANNKGCGGATTATAACTAT |
| RBD_F1-A87-NNK | CAGACCGGCAAAGTGNNKGATTATAACTATAAA |
| RBD_F1-D88-NNK | ACCGGCAAAGTGGCGNNKTATAACTATAAACTG |
| RBD_F1-Y89-NNK | GGCAAAGTGGCGGATNNKAACTATAAACTGCCG |
| RBD_F1-N90-NNK | AAAGTGGCGGATTATNNKTATAAACTGCCGGAT |
| RBD_F1-Y91-NNK | GTGGCGGATTATAACNNKAAACTGCCGGATGAT |
| RBD_F1-K92-NNK | GCGGATTATAACTATNNKCTGCCGGATGATTTT |
| RBD_F1-L93-NNK | GATTATAACTATAAANNKCCGGATGATTTTATT |
| RBD_F1-P94-NNK | TATAACTATAAACTGNNKGATGATTTTATTGGC |
| RBD_F1-F97-NNK | AAACTGCCGGATGATNNKATTGGCTGCGTGATT |

| | |
|---|---|
| RBD_F1-I98-NNK | CTGCCGGATGATTTTNNKGGCTGCGTGATTGCG |
| RBD_F1-G99-NNK | CCGGATGATTTTATTNNKTGCGTGATTGCGTGG |
| RBD_F1-C100-NNK | GATGATTTTATTGGCNNKGTGATTGCGTGGAAC |
| RBD_F1-V101-NNK | GATTTTATTGGCTGCNNKATTGCGTGGAACGCG |
| RBD_F1-I102-NNK | TTTATTGGCTGCGTGNNKGCGTGGAACGCGAAC |
| RBD_F1-A103-NNK | ATTGGCTGCGTGATTNNKTGGAACGCGAACAAC |
| RBD_F1-W104-NNK | GGCTGCGTGATTGCGNNKAACGCGAACAACCTG |
| RBD_F1-A106-NNK | GTGATTGCGTGGAACNNKAACAACCTGGATAGC |
| RBD_F1-N107-NNK | ATTGCGTGGAACGCGNNKAACCTGGATAGCAAA |
| RBD_F1-D110-NNK | AACGCGAACAACCTGNNKAGCAAAGTGGGCGGC |
| RBD_F1-S111-NNK | GCGAACAACCTGGATNNKAAAGTGGGCGGCAAC |
| RBD_F1-G115-NNK | GATAGCAAAGTGGGCNNKAACTATAACTATCTG |
| RBD_F1-N116-NNK | AGCAAAGTGGGCGGCNNKTATAACTATCTGTAT |
| RBD_F1-Y119-NNK | GGCGGCAACTATAACNNKCTGTATCGCCTGTTT |
| RBD_F1-Y121-NNK | AACTATAACTATCTGNNKCGCCTGTTTCGCAAA |
| RBD_F1-R122-NNK | TATAACTATCTGTATNNKCTGTTTCGCAAAAGC |
| RBD_F1-R125-NNK | CTGTATCGCCTGTTTNNKAAAAGCAACCTGAAA |
| RBD_F1-L129-NNK | TTTCGCAAAAGCAACNNKAAACCGTTTGAACGC |
| RBD_F1-D135-NNK | AAACCGTTTGAACGCNNKATTAGCACCGAAATT |
| RBD_F1-I140-NNK | GATATTAGCACCGAANNKTATCAGGCGGGCAGC |
| RBD_F1-C148-NNK | GCGGGCAGCACCCCGNNKAACGGCGTGGAAGGC |
| RBD_F1-C156-NNK | GTGGAAGGCTTTAACNNKTATTTTCCGCTGCAG |
| RBD_F1-P159-NNK | TTTAACTGCTATTTTNNKCTGCAGAGCTATGGC |
| RBD_F1-L160-NNK | AACTGCTATTTTCCGNNKCAGAGCTATGGCTTT |
| RBD_F1-Y163-NNK | TTTCCGCTGCAGAGCNNKGGCTTTCAGCCGACC |
| RBD_F1-F165-NNK | CTGCAGAGCTATGGCNNKCAGCCGACCAACGGC |
| RBD_F1-N169-NNK | GGCTTTCAGCCGACCNNKGGCGTGGGCTATCAG |
| RBD_F1-G172-NNK | CCGACCAACGGCGTGNNKTATCAGCCGTATCGC |
| RBD_F1-Q174-NNK | AACGGCGTGGGCTATNNKCCGTATCGCGTGGTG |
| RBD_F1-P175-NNK | GGCGTGGGCTATCAGNNKTATCGCGTGGTGGTG |
| RBD_F1-Y176-NNK | GTGGGCTATCAGCCGNNKCGCGTGGTGGTGATT |
| RBD_F1-R177-NNK | GGCTATCAGCCGTATNNKGTGGTGGTGATTAGC |
| RBD_F1-V178-NNK | TATCAGCCGTATCGCNNKGTGGTGATTAGCTTT |
| RBD_F1-V179-NNK | CAGCCGTATCGCGTGNNKGTGATTAGCTTTGAA |
| RBD_F1-V180-NNK | CCGTATCGCGTGGTGNNKATTAGCTTTGAACTG |
| RBD_F1-I181-NNK | TATCGCGTGGTGGTGNNKAGCTTTGAACTGCTG |
| RBD_F1-S182-NNK | CGCGTGGTGGTGATTNNKTTTGAACTGCTGCAT |
| RBD_F1-F183-NNK | GTGGTGGTGATTAGCNNKGAACTGCTGCATGCG |
| RBD_F1-V192-NNK | CATGCGCCGGCGACCNNKTGCGGCCCGAAAAAA |

### Table S3: Library Statistics

| Library | SARS-CoV-2 RBD core mutation library - reference population | | | |
|---|---|---|---|---|
| Sub-Library | **Tile1A** | **Tile2A** | **Tile1B** | **Tile2B** |
| Amino Acids | C336-I430 | G431-V524 | C336-I430 | G431-V524 |
| Total Mutated Positions | 52 | 38 | 52 | 38 |
| Total QC Reads (millions) | 0.57 | 0.78 | 0.74 | 0.33 |
| Wild-Type Sequences | 70.0% | 57.2% | 62.8% | 62.6% |
| Sequences with exactly 1 nonsynonymous mutation | 25.5% | 35.7% | 31.4% | 31.6% |
| Sequences with more than 1 nonsynonymous mutation | 4.5% | 7.1% | 5.8% | 5.8% |
| Coverage of all possible single nonsynonymous mutations | 72.6% (755/1040) | 75.1% (571/760) | 74.0% (770/1040) | 68.9% (524/760) |
| Coverage of all possible single nonsynonymous mutations (total) | 73.7% (1326/1800) | | 71.9% (1294/1800) | |

| Library | SARS-CoV-2 RBD core mutation library - 1000 U/ml chymotrypsin treatment | | | |
|---|---|---|---|---|
| Sub-Library | **Tile1A** | **Tile2A** | **Tile1B** | **Tile2B** |
| Amino Acids | C336-I430 | G431-V524 | C336-I430 | G431-V524 |
| Total Mutated Positions | 52 | 38 | 52 | 38 |
| Total QC Reads (millions) | 0.38 | 0.76 | 0.44 | 0.64 |
| Wild-Type Sequences | 74.2% | 63.2% | 68.5% | 70.0% |
| Sequences with exactly 1 nonsynonymous mutation | 21.1% | 29.1% | 25.7% | 24.2% |
| Sequences with more than 1 nonsynonymous mutation | 4.7% | 7.7% | 5.8% | 5.8% |
| Coverage of all possible single nonsynonymous mutations | 74.9% (779/1040) | 81.4% (619/760) | 78.7% (818/1040) | 76.3% (580/760) |
| Coverage of all possible single nonsynonymous mutations (total) | 77.7% (1398/1800) | | 77.7% (1398/1800) | |

| Library | SARS-CoV-2 RBD core mutation library - 2000 U/ml chymotrypsin treatment | | | |
|---|---|---|---|---|
| Sub-Library | **Tile1A** | **Tile2A** | **Tile1B** | **Tile2B** |
| Amino Acids | C336-I430 | G431-V524 | C336-I430 | G431-V524 |
| Total Mutated Positions | 52 | 38 | 52 | 38 |
| Total QC Reads (millions) | 0.37 | 0.35 | 0.74 | 0.59 |
| Wild-Type Sequences | 77.2% | 66.6% | 70.2% | 72.4% |
| Sequences with exactly 1 nonsynonymous mutation | 17.9% | 24.7% | 24.1% | 18.6% |
| Sequences with more than 1 nonsynonymous mutation | 4.9% | 8.8% | 5.7% | 9.0% |
| Coverage of all possible single nonsynonymous mutations | 74.2% (772/1040) | 77.8% (591/760) | 79.9% (831/1040) | 74.3% (565/760) |
| Coverage of all possible single nonsynonymous mutations (total) | 75.7% (1363/1800) | | 77.6% (1396/1800) | |

| Library | SARS-CoV-2 RBD core mutation library - 4000 U/ml chymotrypsin treatment | | | |
|---|---|---|---|---|
| Sub-Library | **Tile1A** | **Tile2A** | **Tile1B** | **Tile2B** |
| Amino Acids | C336-I430 | G431-V524 | C336-I430 | G431-V524 |
| Total Mutated Positions | 52 | 38 | 52 | 38 |
| Total QC Reads (millions) | 0.17 | 0.55 | 0.53 | 0.33 |
| Wild-Type Sequences | 78.3% | 65.7% | 65.5% | 64.5% |
| Sequences with exactly 1 nonsynonymous mutation | 16.3% | 26.6% | 28.1% | 20.4% |
| Sequences with more than 1 nonsynonymous mutation | 5.5% | 7.7% | 6.4% | 15.1% |
| Coverage of all possible single nonsynonymous mutations | 73.6% (765/1040) | 77.6% (590/760) | 75.5% (785/1040) | 72.1% (548/760) |
| Coverage of all possible single nonsynonymous mutations (total) | 75.3% (1355/1800) | | 74.1% (1333/1800) | |

#### Figure S1

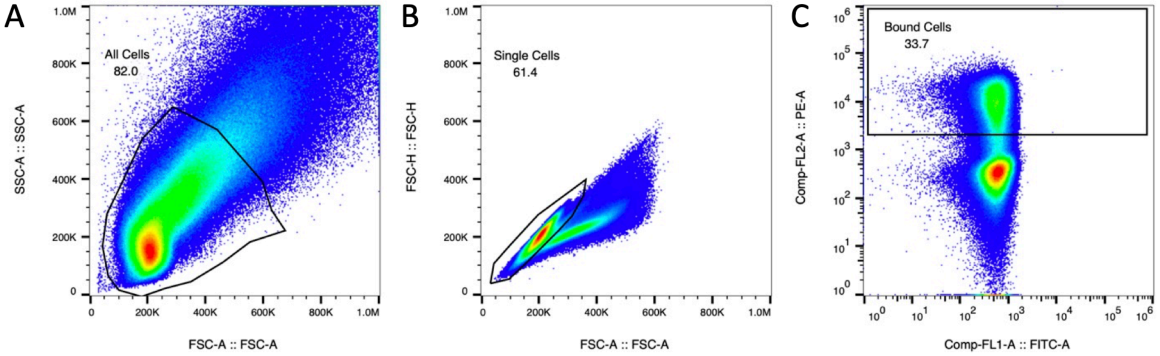

**Figure S1:** ***Fluorescence-Activated Cell Sorter gating for deep mutational scanning screen*. A.** SSC-A/FSC-A gate to discriminate cells from other noise. **B.** FSC-H/FSC-A gate to select only single cells. **C.** PE-A/FITC-A gate to sort cells by binding affinity to ACE2. Bound cell gate collects all cells with a PE signal over 2000.

#### Figure S2

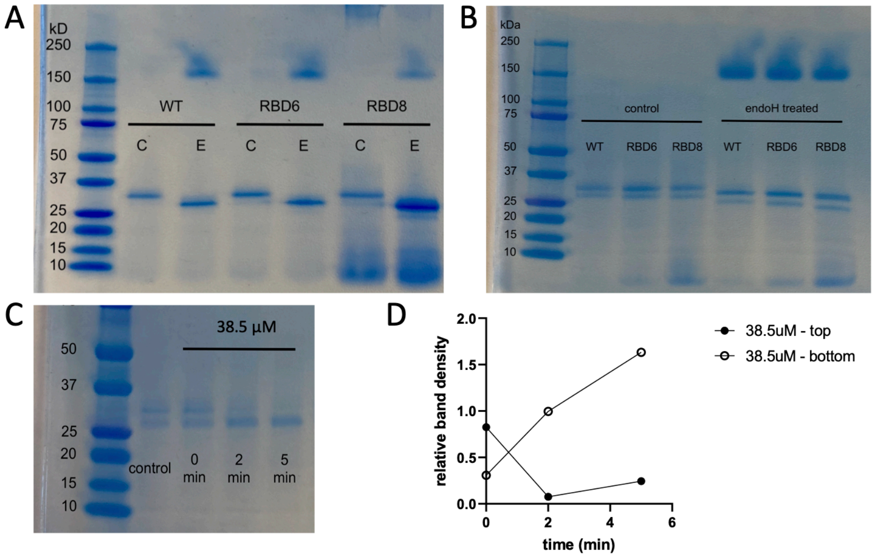

**S2:** ***Degradation of purified RBD protein at 4°C over time can be recapitulated by proteolytic degradation*. A.** SDS-PAGE gel of RBD proteins immediately after Ni-NTA column purification. <ALISON DEFINE “C” LANES>. “E” lanes are treated with Endo H to removed N-linked glycan compared to control “C” lanes without Endo H treatment. **B.** SDS-PAGE gel of RBD proteins run 27 days after purification. **C.** RBD6 protein treated with 38.5 µM or  at room temperature, with the reaction quenched with EDTA immediately (0 min), after 2 min, or after 5 min. **D.** Quantification of gel band intensity from panel C after 38.5 µM (circles) thermolysin treatment. Closed symbols indicate upper band intensity, open circles of lower band intensity. Band intensity is relative to the higher molecular weight band for the control sample.
